# Axonal transcriptome reveals upregulation of PLK1 as a protective mechanism in response to increased DNA damage in FUS^P525L^ spinal motor neurons

**DOI:** 10.1101/2024.11.20.624439

**Authors:** Vitaly Zimyanin, Banaja P. Dash, Dajana Großmann, Theresa Simolka, Hannes Glaß, Riya Verma, Vivek Khatri, Christopher Deppmann, Eli Zunder, Stefanie Redemann, Andreas Hermann

**Affiliations:** Department of Molecular Physiology and Biological Physics, School of Medicine, University of Virginia, Charlottesville, VA, 22903, USA; Center for Membrane and Cell Physiology, School of Medicine, University of Virginia, Charlottesville, VA, 22903, USA; Translational Neurodegeneration Section „Albrecht Kossel“, Department of Neurology, University Medical Centre Rostock, University of Rostock, Rostock, Germany; Department of Biology, Graduate School of Arts and Sciences, University of Virginia, Charlottesville, VA, 22902, USA; College of Dental Medicine, Columbia University, New York, NY, 10032, USA; Department of Biomedical Engineering, School of Medicine, University of Virginia, Charlottesville, VA, 22902 USA; Department of Cell Biology, School of Medicine, University of Virginia, Charlottesville, VA, 22903, USA; Deutsches Zentrum für Neurodegenerative Erkrankungen (DZNE) Rostock/Greifswald, 18147 Rostock, Germany; Center for Transdisciplinary Neurosciences Rostock (CTNR), University Medical Center Rostock, Rostock, Germany

**Author notes:** Correspondence: Prof. Dr. Dr. Andreas Hermann, Schilling Professor for Translational Neurodegeneration, Translational Neurodegeneration Section “Albrecht Kossel”, Dept. Neurology, University Medical Center Rostock, Gehlsheimer Straße 20, 18147 Rostock, Germany, (A.H.). contributed equally (Vitaly Zimyanin, Banaja P. Dash).

**Keywords:** amyotrophic lateral sclerosis, axonal transcriptome, PLK1, cell cycle, axon degeneration, induced pluripotent stem cells, RNA sequencing, differentially expressed genes, DNA damage

## Abstract

Mutations in the gene *FUSED IN SARCOMA* (*FUS*) are among the most frequently occurring genetic forms of amyotrophic lateral sclerosis (ALS). Early pathogenesis of *FUS*-ALS involves impaired DNA damage response and axonal degeneration. However, it is still poorly understood how these gene mutations lead to selective spinal motor neuron (MN) degeneration and how nuclear and axonal phenotypes are linked. To specifically address this, we applied a compartment specific RNA-sequencing approach using microfluidic chambers to generate axonal as well as somatodendritic compartment-specific profiles from isogenic induced pluripotent stem cells (iPSCs)-derived MNs. We demonstrate high purity of axonal and soma fractions and show that the axonal transcriptome is unique and distinct from that of somas including significantly fewer number of transcripts. Functional enrichment analysis revealed that differentially expressed genes (DEGs) in axons were mainly enriched in key pathways like RNA metabolism and DNA damage, complementing our knowledge of early phenotypes in ALS pathogenesis and known functions of FUS. In addition, we demonstrate a strong enrichment for cell cycle associated genes including significant upregulation of polo-like kinase 1 (PLK1) in FUS^P525L^ mutant MNs. PLK1 was increased upon DNA damage induction and PLK1 inhibition further increased the number of DNA damage foci in etoposide-treated cells, an effect that was diminished in case of *FUS* mutant MNs. In contrast, inhibition of PLK1 increased late apoptotic or necrosis-induced neuronal cell death in mutant neurons. Taken together, our findings provide insights into compartment-specific transcriptomics in human *FUS*-ALS MNs and we propose that specific upregulation of PLK1 might represent an early event in the pathogenesis of ALS, possibly modulating DNA damage response and other associated pathways.

## Introduction

Amyotrophic lateral sclerosis (ALS) is a severe and incurable disease that is characterized by degeneration of upper and lower spinal motor neurons (MNs) usually starting by denervation at the distal axons (*1*). The identification of a number of genetic factors associated with ALS and the ability to model this disease in various cell cultures and *in vivo* model systems have contributed enormously to our understanding of pathogenic mechanisms underlying the disease (*2, 3*). While ∼90% of ALS patients suffer from apparently sporadic disease, ∼10% show classical Mendelian heritability with mutations found in over 40 genes (*3*). Mutation of the *FUSED IN SARCOMA (FUS)* gene belong to one of the most common forms of familial ALS and are particularly prevalent in young patients (*4*).

*FUS* encodes a primarily nuclear protein that contains RNA binding- and low-complexity- domains (LCD) and has been implicated in regulating transcription, DNA damage, RNA splicing, nucleocytoplasmic and axonal trafficking, phase-separation dynamics of ribonucleoprotein assemblies, and metabolism (*5*). Many of the disease causing mutations are located within nuclear localization sequence (NLS) (*6*), and cytoplasmic mislocalization of FUS is a pathophysiological hallmark of *FUS*-ALS (*7*). FUS binding was mapped to multiple sites on chromosomes and more than a thousand long coding mRNAs, where it was shown to regulate their splicing (*8–12*). This ability of FUS to regulate many identified binding targets is likely to be central to ALS progression and development. However, the detailed contribution of such potentially global misregulation, as well as hierarchy of contributing pathways in disease progression remains unclear. Some of the downstream defects in ALS clearly converge on or are triggered by the formation of toxic proteins or ribonucleoprotein (RNP) aggregates (*13*). In this respect ALS is similar to other neurodegenerative diseases, where multiple cellular mechanisms drive the vulnerability of their specific subsets of neuronal cells (*13*).

Previous studies have shown that changes in mRNA and protein levels can occur in specific neuronal compartments, such as distal axons, and thus these changes could provide insight into the underlying mechanisms of ALS (*14, 15*). Interestingly, the pathogenesis of ALS is often referred to as axonopathy, with particular vulnerability of distal axons/neuromuscular junctions and muscle denervation being the first symptoms of the disease (*16*). We and others have shown that this is particularly true for *FUS*-ALS (*17–19*). Induced pluripotent stem cell (iPSCs)-derived MNs of *FUS*-ALS exhibited distal axonopathy with depolarized distal axonal mitochondria prior to a dying-back and nerve cell loss. *FUS*-ALS MNs also accumulated DNA damage, and DNA damage induction was sufficient to induce distal axonal mitochondrial depolarization (*17, 20*). While axonal ATP levels were lower in general in iPSC-derived MNs, no axonal compartment specific differences in ATP levels in MNs with *FUS*-ALS mutations have been yet detected (*21*), nor have reductions in ATP consumption been observed (*22, 23*). Thus, the exact mechanisms by which nuclear DNA damage affects axonal mitochondrial integrity are so far not understood.

Recent advances in high-throughput transcriptomic profiling based on next-generation RNA-sequencing (RNA-seq) approaches have provided insights to better understand gene expression analysis/mechanisms of ALS pathogenesis in healthy and diseased conditions (*24–27*). However, the transcriptome profiles often show surprisingly little overlap in what should be comparable or at least converging disease models, partially due to genetic background differences and purity of cell populations (*28*). These global quantifications also lack information about local changes of the transcriptome in different cellular compartments. RNA-seq methods aimed at purifying the axonal transcriptome have been used as a novel tool to precisely quantify local changes of mRNA composition and its modulation in the axonal fractions of neurons derived from primary mouse MNs and primary mouse dorsal root ganglia (DRG) (*29–34*). More recently, Nijssen and Aguila *et al*. developed an elegant Axon-seq approach, a robust method incorporating microfluidics with sensitivity similar to single-cell sequencing, and compared transcriptome changes in mouse and human stem cell-derived spinal motor axons in healthy controls and mutant lines (*35*). To the best of our knowledge, only very few studies have paid attention to the important roles of distinct mRNA expression profiles of the axon and soma by combining microfluidic approaches and iPSC-derived MNs from ALS patients and specifically for FUS-related models (*35, 36*). Therefore, to investigate transcripts and molecular pathways critical for axonal function and degeneration, we developed a microfluidic-based system that enabled us to quantify *FUS* mutant dependent local changes to the axonal transcriptome. Using previously generated isogenic CRISPR/Cas9 edited iPSC-lines harboring *FUS^P525L^* mutation (*17, 37, 38*), coupled with microfluidic approaches we here profiled and compared transcriptomes in somatodendritic (SD) and axonal compartments. We aimed to gain insight into the local transcriptomic alterations (soma versus axon) leading to differentially expressed genes (DEGs) and pathways in the *FUS* mutant background, and into the functional consequences of these changes in MNs. Differential analysis of mRNA expression in SD and axon compartments strengthened and highlighted a number of potential pathways including DNA damage, mRNA splicing and mitochondrial functions. Further, we show altered DNA damage response as a functional consequence of cell cycle related gene polo-like kinase 1 (PLK1) upregulation in mutant MNs, and this activation has important implications on survival of *FUS*-ALS MNs.

## Results

### Compartmentalized RNA-seq of iPSC-derived FUS^WT^ and FUS^P525L^ MNs

The P525L mutation in *FUS* has been associated with an early onset and an aggressive form of ALS. In our lab, we have previously generated and characterized an isogenic line carrying this single nucleotide substitution (hereafter FUS^WT^ and FUS^P525L^). The used small molecule based published protocol allows a high yield of cells that express all major motor neuronal cell markers and demonstrate disease-related phenotypical changes at well-defined time points (*17, 21, 39*) (Figure 1A). We quantified FUS protein localization in our line by staining for FUS protein and detected FUS protein mislocalization to the cytoplasm in FUS^P525L^ mutant lines (Figure 1B-C). We used this cell line to perform a comprehensive subcellular transcriptomic analysis to identify changes that occur in unstressed mutant cells. Additionally, since ALS disease etiology and previous results have shown that some early pre-symptomatic changes occur in axonal compartment, we decided to utilize microfluidic devices to isolate RNA from distal compartments of axons (Figure 1D). We chose the day 14 *in vitro* (DIV14) stage of MN maturation as the analysis time point since at this stage axonal morphology is still intact whereas distal axonal organelle trafficking phenotypes can already be detected in the *FUS*-ALS MNs (Figure 1A) (*21, 40*). We then used resulting RNA libraries, which were polyA-amplified for axonal samples, and Illumina sequencing to compare the transcriptome of control and FUS^P525L^ mutant MNs.

**Figure 1.**
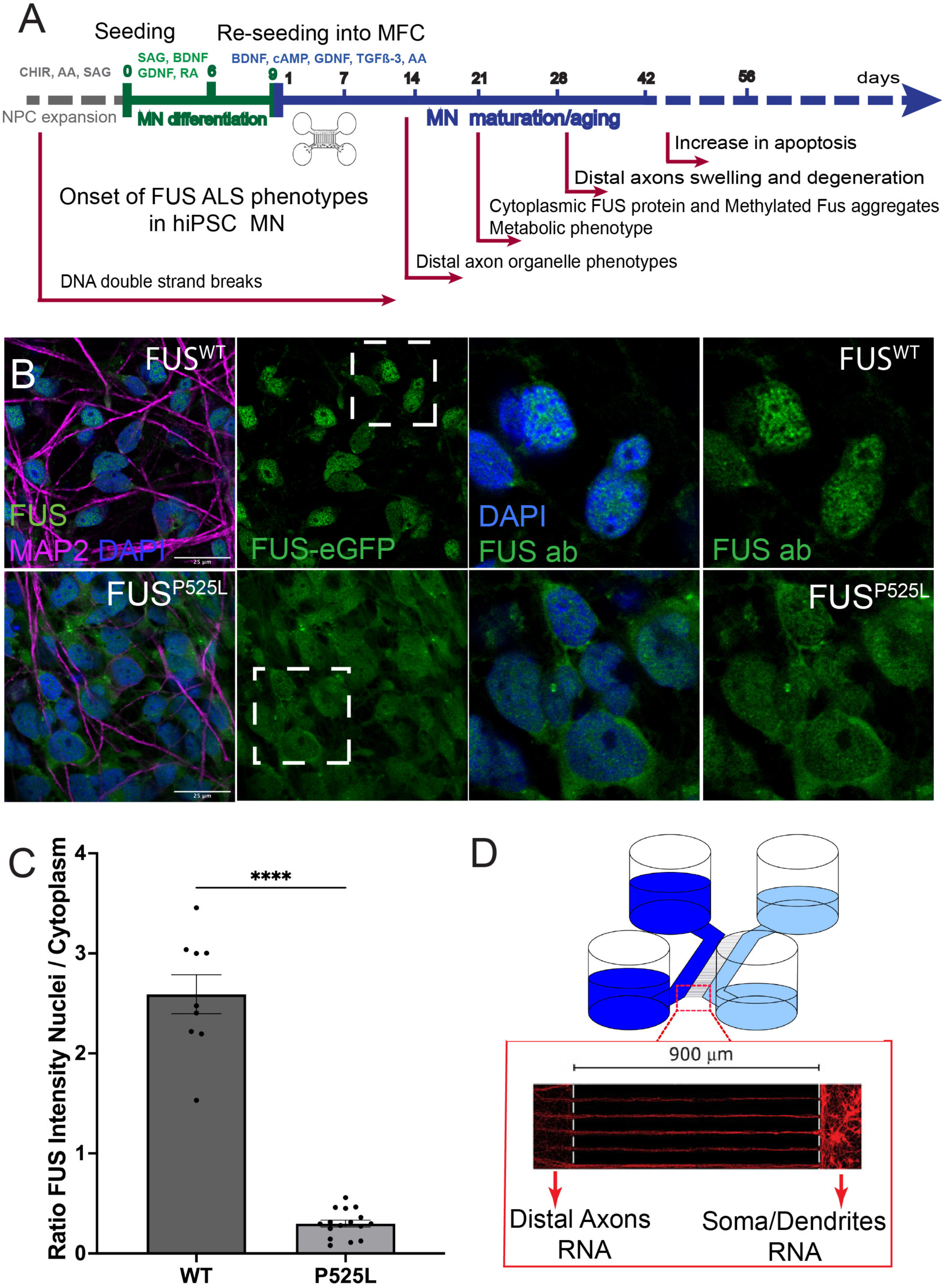
Compartmentalized transcriptomics of FUS^P525L^ MNs. (A) Timeline and schematic outline of the MN differentiation protocol starting from neural precursor cells and undergoing sequential differentiation and maturation. Time points for seeding into MFC and onset of previously reported *FUS*-ALS-related phenotypes are indicated under the timeline. (B) Mature cultured MNs showing mis-localization of mutant FUS^P525L^ protein from the nucleus to the cytoplasm, as seen by fluorescence imaging of *FUS*-eGFP tagged isoforms (left) or by immunostaining with FUS antibody (right) and quantified in (C). (C) Schematic of a microfluidic device and polarized neuron growth used to purify separate RNA from SD and distal axonal compartments.

Our RNA-seq workflow encompassed read quality control (QC), genome mapping, and gene quantification (see Methods). RNA-seq libraries were analysed from an average depth of ∼21-30 million fragments 75 bp single-end reads of good quality (DV200 ≥ 60%) and low amount of rRNA reads (1.1% ± 0.25%). Sequencing reads were then processed to generate normalized counts and the libraries compared using principal component analysis (PCA), indicating SD samples clearly separated from axon samples (Supplementary Figure S1A) and is in agreement with Pearson and Spearman’s and unsupervised hierarchical clustering analysis. Pearson and Spearman’s rank correlation coefficient clustering (Supplementary Figure S1B-E) revealed similar separation of SD from all other samples. Among the whole dataset, the SD samples exhibited an average of 2,38x10^4^ (± 264) detectable genes, whereas the axon samples demonstrated remarkably less with an average of 1,3x10^4^ (± 3251) genes (Supplementary Table S1). Interestingly, DEGs analysis showed as minimal overlap between both compartments with 97 upregulated genes and 69 downregulated genes from SDs (n = 166, false discovery rate (FDR) value ≤ 0.05 and 123 upregulated genes and 14 downregulated genes (n = 137) from the axon compartment, selected at the same threshold (Figure 2A-C). The unsupervised hierarchical clustering heatmap analysis of the most prevalent DEGs revealed significant differences in expression patterns between the SD and axonal compartments, highlighting those top 50 genes with the highest mean expression strength and highly variable genes across SD and axon samples (FUS^P525L^ versus FUS^WT^ MNs) were nicely grouped into separate clusters (Figure 2D-E, Supplementary Figure S1F-G). A full list of DEG read values is given in (Supplementary Table S1). In summary, this shows that our experimental pipeline enables RNA-seq of both SD and pure distal axonal preparations of human iPSC-derived MNs with a high selectivity and specificity.

**Figure 2.**
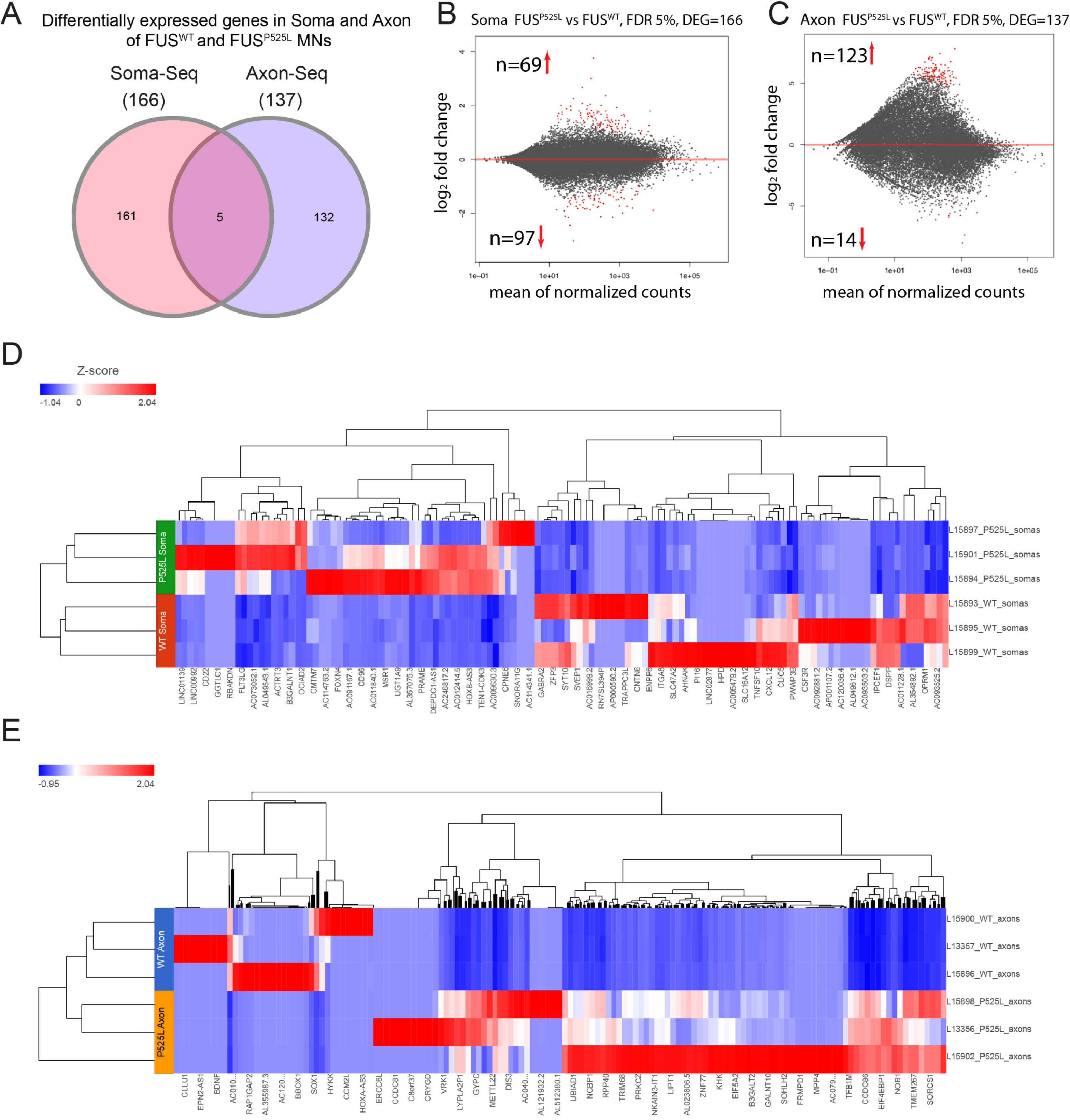
RNA-seq analysis identifies distinct compartment-specific DEGs. (A) Venn diagram showing number of detected differentially expressed genes in the axons and SDs of the FUS^WT^ and FUS^P525L^ MNs (log2FC ≥ 1 or log2FC ≤ −1, adjusted *p*-value ≤ 0.05). (B-C) MA scatter plots depict significantly changed (adjusted *p*-value ≤ 0.05) DEGs (red dots) with FDR 5% for SD (B) and axons (C) displaying the mean expression (x-axis) and the expression change (y-axis) between conditions. (D-E) The heatmap showing the differential expression of significant (adjusted *p*-value ≤ 0.05, log2FC ≥1) mRNAs in the soma (D) or axon (E) compartment of iPSC derived FUSWT and FUSP525L MNs. Heatmaps were generated in *Partek*™ *Flow*™ software, v11.0 using normalized (separate Z-score calculated per gene) expression values (DESeq2) across all samples. Each row in the heatmap represents a sample, and each column represents a mRNA and the color scale at the left of the heatmap represents the Z-score ranging from blue (low expression) to red (high expression), respectively.

### Extracellular secretion and extracellular matrix disassembly pathways are important in all distal axonal compartments

To additionally verify the validity of subcellular compartment-specific transcriptome of iPSC-derived MNs in general, we compared DEGs between SD and axonal compartments of the FUS^WT^ healthy control samples only (Supplementary Table S2). DEG analysis identified 161 genes that were higher expressed in control axons as compared to SD. Enrichment of the latter genes using the gene ontology (GO; biological process (BP), molecular function (MF), and cellular component (CC) and pathway terms detected inflammatory response, endopeptidase activity, extracellular matrix, extracellular matrix organization (ECM), and activation of metalloproteases pathways/processes, some of which could be important for the correct function and growth of distal growing axons (Supplementary Figure S2A-C). DEG analysis also identified a large number of lower expressed transcripts in axons (2482 genes), and these genes were associated with DNA/RNA metabolic processes, nucleic acid catalytic activity, postsynaptic membrane, DNA repair, cell cycle, and PLK1 pathway, among others, most of which are not usually associated with the axonal functions and were expected to be lower than in the somatodendritic compartment (Supplementary Figure S2A-C). GO and pathway enrichment results are summarized in Supplementary Table S3. To further identify the functional alterations in specific molecular phenotypes of FUS^WT^ motor axons, we performed gene set enrichment analysis (GSEA) (*41*) using the GO term and KEGG pathway databases. Particularly, in axons (versus soma) we found consistent increase of gene sets enrichment in cell-matrix adhesion, tight junction, and WNT signaling, whereas a large number of gene sets demonstrated significant lower expression of RNA-polymerase based gene transcription, DNA repair, regulation of immune system process, and mitochondrial matrix, respectively (Supplementary Figure S3A-L; Supplementary Table S4).

Interestingly, fewer increased transcripts (25 genes) were detected when comparing axonal and SD transcriptome within FUS^P525L^ MNs. Their enrichment analysis showed some similarity to control enriched terms and pathways, such as secreted peptidase/proteases, peptidoglycan binding, vesicle lumen and ECM disassembly, respectively (Supplementary Figure S2D-F). On the other hand, lower expressed genes in axons compared to soma (n=332) included a notable prevalence of DEGs linked to cell cycle regulation and DNA damage response. Distinct for FUS^P525L^ MN axon-specific pathways were glycogen metabolism, WNT signaling and organelle biogenesis (Supplementary Figure S2D-F). GO and pathway enrichment results are summarized in Supplementary Table S3. Furthermore, GSEA also confirmed significant enrichment with increased expression of mRNA splicing, various types of N-glycan biosynthesis, TCA cycle, and WNT signaling gene sets among others (Supplementary Figure S4A-H, Supplementary Table S4). In axons (versus soma), GSEA of lipid metabolic process, trans-Golgi network, and MAPK signaling pathway gene sets revealed significant reduced axonal expression in FUS^P525L^ MNs, which can contribute to the dysregulation of the fine tuning between metabolism and cell survival during development (Supplementary Figure S4A-H, Supplementary Table S4). Taken together, these data show clear functional differences between transcribed RNA that is present in SD and distal axonal compartments of the same cells. Such differences could be critical for the correct function of axons, highlighting the axonal role of extracellular secretion and ECM that could be perturbed in diseased MNs.

### Cross-comparison to published axonal transcriptomes highlights processes characteristic for distal axons

Axonal-specific transcriptome is a novel way to look at the compartment specific disease mechanisms in ALS. Very few studies have analyzed axonal transcriptomes from human and mouse stem cell-derived MNs (*29, 35*). To understand which mRNAs are specific to human motor axons and if there was conservation in the intrinsic transcriptomic signature/pathways/DEGs central to MN axons, we compared our axonal-seq datasets with the single published axon datasets derived from iPSC MNs (Nijssen, Aguila et al. 2018). In this comparison of WT MNs, pathway analysis on the overlapping genes (453 out of 5445 transcripts) revealed significant transcripts critical for axonal functions including nervous system development, axon guidance and RNA metabolism, including splicing and translation (Supplementary Figure S2G; Supplementary Table S5). To further investigate if our findings were relevant more broadly across mouse MNs, we compared our axon datasets with previously published axonal-seq datasets derived from primary mouse MNs and mouse embryonic stem cell (mESC)-derived MNs (*29, 35*). This analysis identified an overlap of 60 common genes, several of which were mainly enriched in Rho GTPase and Notch signaling, as well as somewhat unexpectedly cell cycle and PLK1 pathways (Supplementary Figure S2H; Supplementary Table S5). In summary, our cross-comparison analysis of MNs derived from human and mouse stem cells or primary MNs, revealed a broadly shared axonal transcriptional signature including known critical axonal related functions, such as RNA metabolism, splicing and translation, as well as ones not usually associated with axons such as PLK1 and cell cycle pathways. These processes, however could be critical for distal axon maintenance in disease MNs.

### Somatodendritic changes upon FUS mutation include WNT signaling, mitochondrial functions and ECM- and synapse-related functions

For our core analysis we looked at the differences between transcriptomes of FUS^P525L^ and FUS^WT^ MNs. Remarkably, our analysis showed only a small overlap of DEGs between SD and axonal compartments (Figure 2A), suggesting compartment-specific transcriptional misregulation in the case of *FUS*-ALS. In the soma of FUS^P525L^ MNs upregulated genes were enriched in the regulation of WNT signaling pathway and WNT-protein binding activity in the biological process and molecular function ontology categories, whereas inner mitochondrial membrane and Golgi compartments were identified as major cell compartment (Figure 3A). Among the DEGs downregulated in FUS^P525L^ MN somas were several key transcripts critical for neuron development and synapse functions and were enriched in the important biological processes including synaptic signaling, chemical synaptic transmission and axon development. In the molecular function and cell component GO categories, enriched terms were associated with neuropeptide receptor activity, ECM binding, and dendrite (Figure 3B). Furthermore, pathways of upregulated genes were primarily involved in WNT signaling, GPCR signaling and pathways of neurodegeneration (Figure 3C). The downregulated enriched categories comprised items related to neuronal developmental functions, which included neuronal differentiation, ECM and pathways in peptide ligand-binding receptors (Figure 3C). The complete GO and pathway analysis results can be found in Supplementary Table S6.

**Figure 3.**
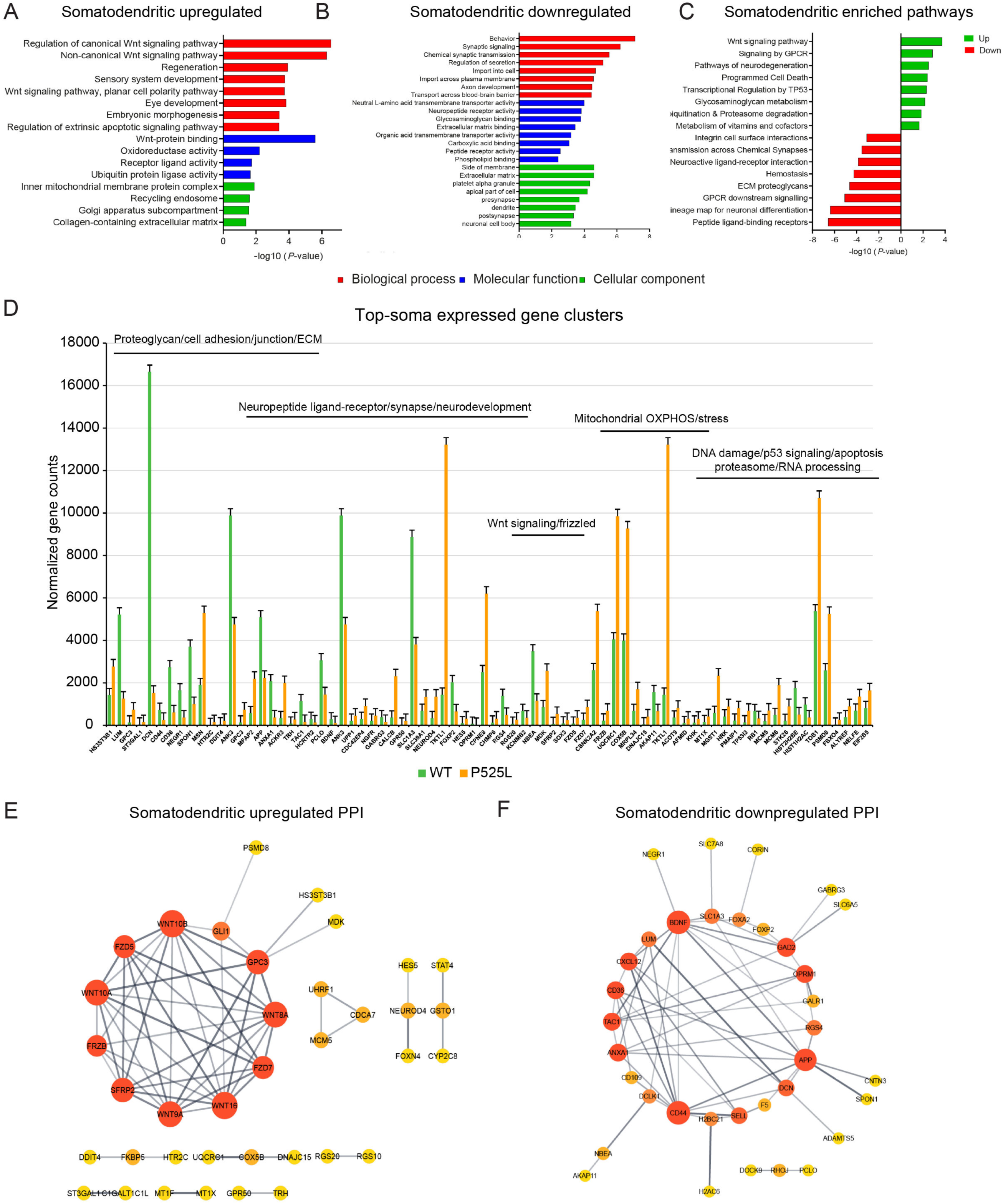
RNA-seq revealed GO, pathway and network analysis of DEGs in SD compartment of FUS^P525L^ MNs. (A-C) Functional analysis of SD-enriched genes identified a variety of (A) upregulated and (B) downregulated GO terms and (C) enriched pathways associated with FUS^P525L^ MNs neurodegeneration (*p* ≤ 0.05). Gene ontology (GO) term enrichment analysis of differential expression genes ranked the most enriched gene group changes in the SD compartment of FUSP^525L^ mutant versus FUS^WT^ healthy control MNs. The enrichment analysis was performed by Metascape and significant terms/pathways were listed with a *p*-value ≤ 0.05. The GO terms were divided into three categories, including biological process (BP, red), molecular function (MF, blue), and cellular component (CC, green), respectively. See also Supplementary Table S6. (D) Bar graph showing expression levels of the top transcripts in FUS^P525L^ and FUS^WT^ SD related to distinct gene clusters, including ECM, synapse/neuronal functions, WNT signaling, mitochondrial OXPHOS and DNA damage response. All of the genes plotted are significantly enriched in the SD compartment of FUS^P525L^ and FUS^WT^ MNs (*p* ≤ 0.05). *p* values were derived from the DESeq2 differential expression and were adjusted for multiple testing. Data are represented as mean ± SEM. (E-F) STRING analysis of differentially regulated genes involved in protein-protein interactions (PPI) between (E) upregulated and (F) downregulated SD-specific genes showed distinct gene clusters. In the PPI network, each node represents a protein/DEG, and each edge represents an interaction. Node size indicates node degree value and edge color represents STRING enrichment score. Darker/thicker edges mean higher/stronger STRING score. The color of a node depends on its degree values (hubs); the darker the color, the higher the connectivity. Nodes depicted in darker color and with large shape represent their importance to the network. The rank and score of these key genes/clusters were identified with high degree of connectivity within the network (topological parameters), with higher scores in red, middle-range scores in orange, and lower scores in yellow. STRING enrichment analysis for protein-associations of the mapped upregulated and downregulated DEGs in SD datasets showed distinct gene clusters involved in WNT signaling pathway and neuronal- and synapse-related functions. The STRING database was used to establish functional connections among the known and predicted proteins using annotated DEGs as query for SD-specific (FUS^P525L^ versus FUS^WT^ MNs) interaction networks (confidence score ≥ 4) and the generated networks were analyzed as well as visualized using Cytoscape v3.10.1.

GSEA analysis on SD-enriched transcripts in FUS^P525L^ showed significant upregulation of genes involved in regulation of apoptotic processes and apoptotic mitochondrial changes, whereas downregulated genes were predominantly involved in nervous system development, proteolysis involved in cellular protein catabolic processes and trans-Golgi network terms (Supplementary Figure S5A-H; Supplementary Table S7). Interestingly, expression levels of the highly expressed transcripts in both FUS^P525L^ and FUS^WT^ SD were related to clusters such as ECM/proteoglycan, neurodevelopmental, WNT signaling, mitochondrial OXPHOS and DNA damage/p53 functions, indicating both nuclear- and mitochondrial-encoded transcripts were distinctly enriched in SD and that most of the abundant transcripts have major involvement in the ECM/proteoglycan and the neuronal development, and mitochondrial energy production related functions or processes (Figure 3D). In addition, we used the STRING v12.0 database to map protein–protein interaction (PPI) networks of the mRNA products identified in SD-specific transcripts (148 matched out of 166 identified DEGs) (*42, 43*). With the medium confidence level, the upregulated PPI network was constructed with an average node degree of 1.32 and the PPI enrichment *P* = 4.96 × 10^−9^. The network with the upregulated genes (FZD7, FZD5, SFRP2, and FRZB) was significantly enriched in WNT signaling pathway (Figure 3E; Supplementary Table 8), while the downregulated network (average node degree of 2.03 and the PPI enrichment *P* = 1.58 × 10^−11^) was notably enriched in ECM related components (such as proteoglycan, cell adhesion/junction) and synaptic functions (Figure 3F; Supplementary Table S8). Taken together, these findings indicate that SD of FUS^P525L^ mutant MNs increase the expression of genes regulating neuronal growth/developmental-related functions while concomitantly downregulating genes involved with ECM- and synapse-related functions.

### Functional enrichment and PPI analysis of axon DEGs centre on PLK1 pathway, mitochondrial gene expression and regulation of inflammation

Changes in the axonal transcriptome of FUS^P525L^ MNs were of special interest in our dataset, as the distal part of the MNs displays some of the earliest visible phenotypes in ALS patients and study model systems. The first obvious difference to the SD compartment was the abundance of DEGs being upregulated in the case of the *FUS* mutant, while more DEGs were downregulated rather than upregulated in the SD compartment (Figure 2B). Using GO analysis, we observed that these upregulated genes in FUS^P525L^ mutant MN axons were significantly enriched in mitochondrial transcription and RNA metabolic (ncRNA, tRNA, rRNA) processes, RNA-binding, microtubule (MT) motor related functions and mitochondrial matrix components (Figure 4A), whereas the few downregulated genes were primarily associated with cell proliferation, proteoglycan relevant processes, ion channel activity functions and membrane raft cellular components, implicating aberrant local regulation of distal axonal related processes or functions in the context of FUS^P525L^ mutations (Figure 4B). Pathway analysis on axon enriched genes revealed that those upregulated in FUS^P525L^ MNs were enriched in the polo-like kinase 1 (PLK1) pathway, mitochondrial gene expression and RNA processing pathways. Conversely, genes decreased in expression in FUS^P525L^ MNs were enriched for ECM related functions (Figure 4C), consistent with shared transcriptional alterations between SD and axon compartments. The complete GO and pathway analysis results can be found in Supplementary Table S9.

**Figure 4.**
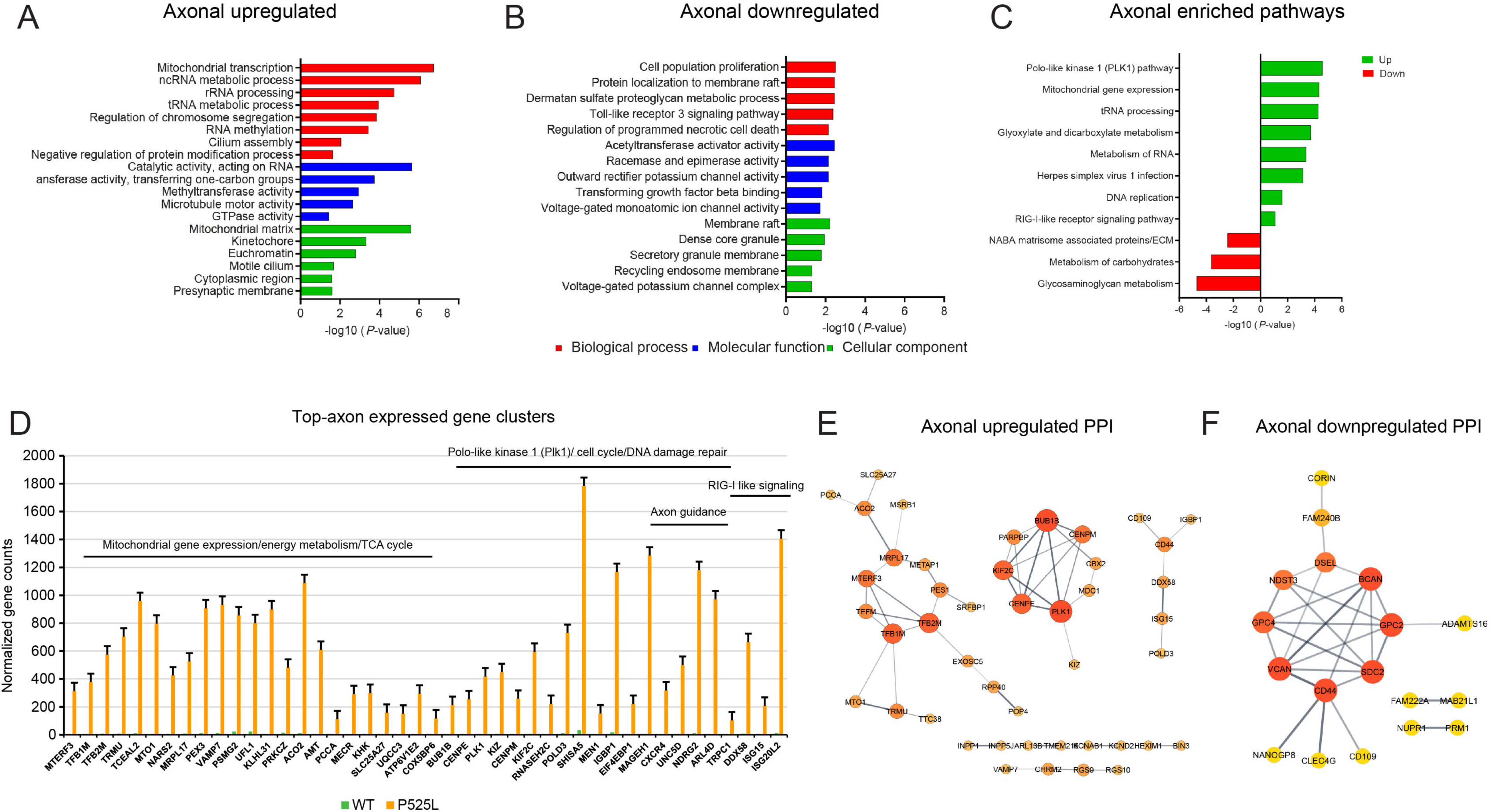
RNA-seq revealed GO, pathway and network analysis of DEGs in axonal compartment of FUS^P525L^ MNs. (A-C) Functional analysis of axonal-enriched genes identified a variety of upregulated (A) and downregulated (B) GO terms and enriched pathways (C) associated with FUS^P525L^ MN neurodegeneration (*p* ≤ 0.05). Gene ontology (GO) term enrichment analysis of differential expression genes ranked the most enriched gene group changes in the axon compartment of FUS^P525L^ mutant versus FUS^WT^ healthy control MNs. The enrichment analysis was performed by Metascape and *p*-value ≤ 0.05 represents the importance of enrichment. Significantly enriched GO terms in biological process (BP), molecular function (MF), and cellular component (CC) groups were displayed in red, blue and green bars, respectively. See also Supplementary Table S9. (D) Bar graph representation of the top transcripts with the highest expression in axons, a majority of which were connected to functional clusters, including mitochondrial transcription/metabolism, PLK1 pathway or DNA damage, axon guidance and RIG-I signaling pathways. All the normalized genes plotted are significantly enriched in the axonal compartment of FUS^P525L^ and FUS^WT^ MNs (*p* ≤ 0.05). *p* values were derived from the DESeq2 differential expression and were corrected for multiple testing. (mean ± SEM). (E-F) STRING analysis of differentially regulated genes involved in protein-protein interactions (PPI) between (E) upregulated and (F) downregulated axon-specific genes showed distinct gene clusters. In the PPI network, each node represents a protein/DEG, and each edge represents an interaction. Node size indicates node degree value and edge color represents STRING enrichment score. Darker/thicker edges mean higher/stronger STRING score. The color of a node depends on its degree values (hubs); the darker the color, the higher the connectivity. Nodes depicted in darker color and with large shape represent their importance to the network. The rank and score of these key genes/clusters were identified with high degree of connectivity within the network (topological parameters), with red to yellow-colored nodes represents genes with high to low PPI degree scores. STRING enrichment analysis for protein-associations of the mapped upregulated and downregulated DEGs in axon datasets showed separate gene clusters involved in mitochondrial functions, cell cycle/cytoskeletal organization including PLK1 pathway, RIG-I signaling pathway and ECM-associated processes. The STRING database was used to establish functional connections among the known and predicted proteins using annotated DEGs as query for axon-specific (FUS^P525L^ versus FUS^WT^ MNs) interaction networks (confidence score ≥ 4) and the generated networks were analyzed as well as visualized using Cytoscape v3.10.1. The STRING database was used to establish functional connections among the known and predicted proteins using annotated DEGs as query for axon- and soma-specific (*FUS*-ALS versus healthy controls) interaction networks (confidence score ≥ 4) and the generated networks were analyzed as well as visualized using Cytoscape v3.10.1.

GSEA further confirmed that genes involved in mitochondrial RNA/DNA metabolic processes and metabolism as well as DNA damage repair pathways were significantly increased in expression in FUS^P525L^ MN axons, whereas downregulated genes were enriched in several processes/pathways, including regulation of mitochondrial potential, focal adhesion, MAPK and PI3K-Akt signaling pathways (Supplementary Figure S6A-J; Supplementary Table S7). Many of these dysregulated genes have previously been implicated in axon function and in ALS pathology. Notably, the majority of transcripts with an increased expression in axons were significantly enriched in mitochondrial gene expression/energy metabolism, PLK1 related cell cycle events/DNA damage and RIG-I signaling as well as axon guidance clusters (Figure 4D). To verify the functional connectivity of the axon-enriched DEGs, a PPI network was generated based on our upregulated genes (116 matched out of 137 DEGs, average node degree 0.879, PPI enrichment *P* = 7.83 × 10^−2^). Notably, for distal axons we identified three distinct PPI clusters formed by highly connected upregulated protein nodes involved in 1) mitochondrial gene expression/energy metabolism (TEFM, TFB1M, TFB2M, and MTERF3, MRPL17, MTO1, TRMU, ACO2, SLC25AA27), 2) cell cycle-based process/MT cytoskeletal organization (PLK1, KIF2C, BUB1B, CENPM, CENPE, and PARPBP), and 3) RIG-I interferon signaling/inflammation pathways (DDX58, ISG15, CD44, IGBP1, CD109, and POLD3) (Figure 4E; Supplementary Table S8), potential candidates for specific MN axon (dys-) functions. Subsequently, downregulated genes in the PPI network (average node degree 2.45, PPI enrichment *P* = 2.74 × 10^−5^) were connected to proteoglycan binding and ECM related processes (Figure 4F; Supplementary Table S8), confirming their essential role in maintaining the normal functions of the motor axons such as axonal growth/regeneration, myelination and cytoskeletal regulation (*44*). Collectively, our findings indicate that growing axons of FUS^P525L^ MNs are enriched for mRNAs associated with mitochondrial and cell cycle related DNA repair machinery. Upregulation in these pathways may have a critical role in maintaining distal axon energy metabolism and survival/neuronal support as a response to DNA damage pathway activation, an early marker of *FUS*-ALS.

### PLK1 expression is upregulated in FUS^P525L^ MNs

Consistent with our results, mutations in familial ALS causing genes such as *TARDBP*, *SOD1*, and *PFN1* were previously associated with upregulation of cell cycle related processes including DNA damage response, and upregulation of PLK1 (*17, 20, 27, 45–47*). Therefore, we decided to test if PLK1 could be an important key protein and pathway responding to FUS mislocalization in MNs. To confirm that PLK1 transcript levels are increased in FUS^P525L^ MNs we performed qPCR analysis and found a 2.2±0.2 fold increase in PLK1 mRNA expression in FUS^P525L^ MNs (Figure 5A). This result was supported by a 3.4±0.82 - fold-increase in PLK1 protein levels (Figure 5B, C). Immunostaining for PLK1 protein showed a relatively weak uniform staining of the MN cells with a small proportion (6.23%±0.96) of control cells showing stronger PLK1 signal at DIV14 (Figure 5D, E).

**Figure 5.**
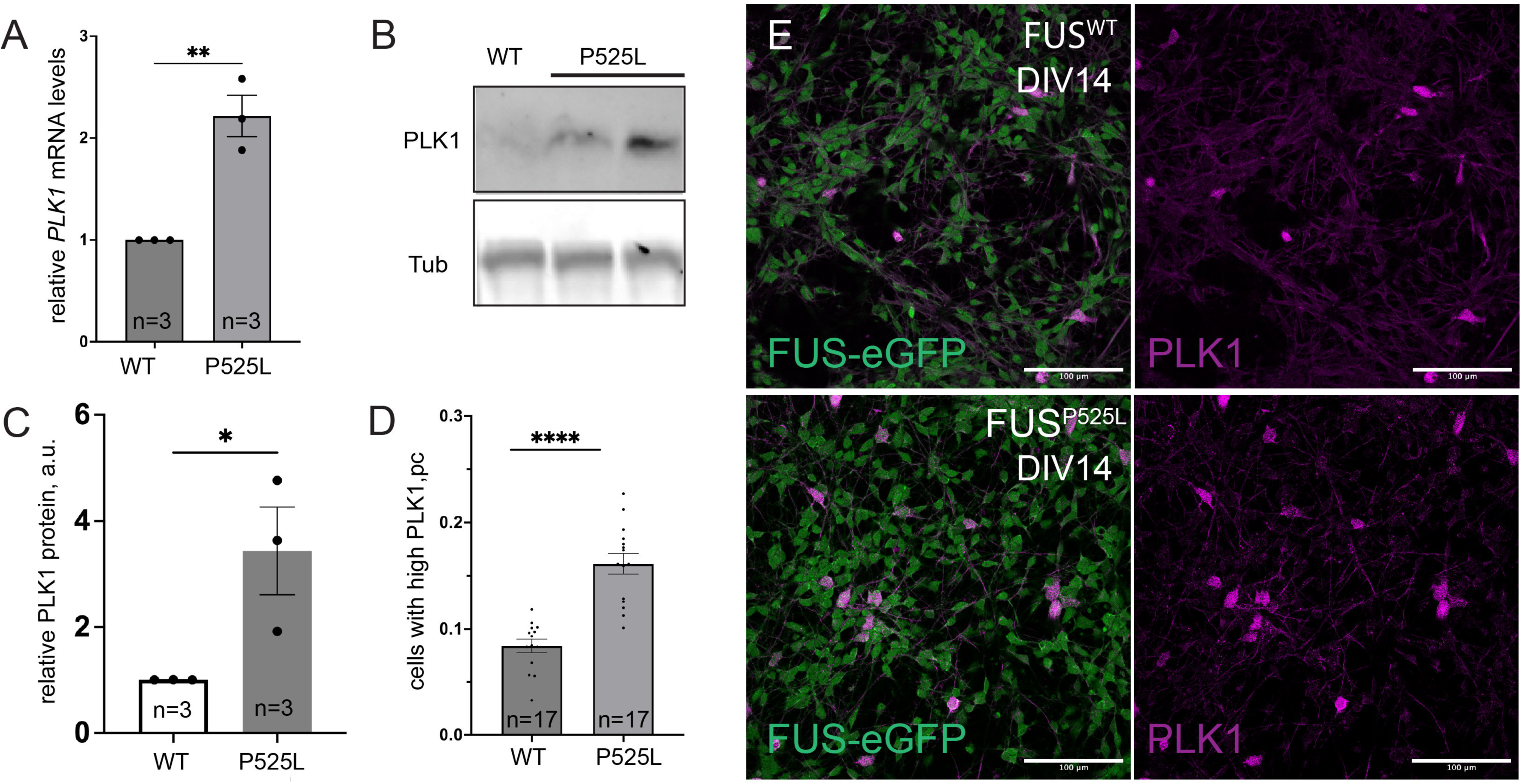
PLK1 is upregulated in FUS^P525L^ MNs. (A) qPCR analysis confirming global relative upregulation of PLK1 mRNA in FUS^P525L^ MN cultures. (B-C) Representative Western blot (B) and quantification (C) of PLK1 protein levels in whole cell lysates of control and FUS^P525L^ MNs (D-E) Immunostaining for PLK1 shows enrichment of PLK1 in a subset of differentiated MN at DIV14 is increased in FUS^P525L^ mutant MN. (D) Quantification of the relative percent of cells showing PLK1 expression in MNs relative to total number of cells at DIV14 and (E) representative images. Significance assessed with unequal variance *t*-test. * *p* ≤ 0.05, ** *p* ≤ 0.01, *** *p* ≤ 0.001, **** *p* ≤ 0.0001.

FUS^P525L^ mutant cells had a significantly higher proportion of such cells (14.23%±1.55). These results indicate that PLK1 activation could be part of the phenotype in *FUS*-ALS MNs.

### FUS^P525L^ MNs do not show cell cycle re-entry but are vulnerable against PLK1 inhibition

The canonical function of PLK1 is required in proliferating cells, during the cell cycle, as well as in cell survival (*48*). We first hypothesized that selective upregulation of PLK1 in the case of *FUS* mutation in MNs is associated with pathological re-entry into the cell cycle, a common feature in a range of neurodegenerative diseases (*49*). To test this hypothesis, we performed cell cycle analysis of neural precursor cells (Supplementary Figure S7A, B and MAP2 positive MNs (Supplementary Figure S8, S9). There was no significant difference in any of the phases of the cell cycle, neither in dividing NPCs nor in post mitotic MNs (Figure 6A) (Supplementary Figure S7-9).

**Figure 6.**
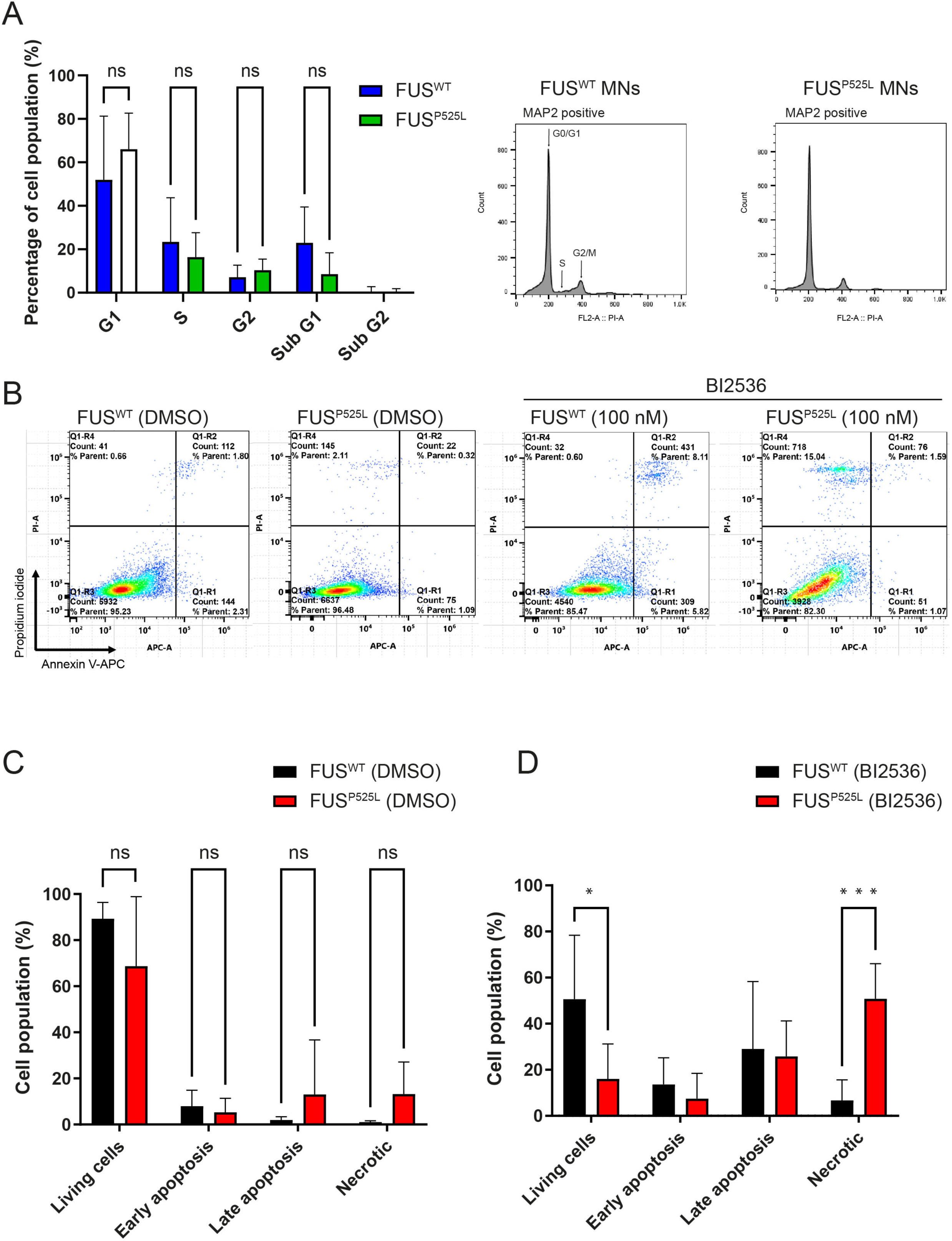
FUS^P525L^ MNs do not show cell cycle re-entry but are vulnerable against PLK1 inhibition. (A) Percentage of cell cycle stages/distribution calculated from the histogram using the Watson pragmatic algorithm in FLowJo showed no significant effect or change on MAP2 positive cells in FUS^P525L^ compared to FUS^WT^ MNs. Cell cycle profiles (right panel) of untreated MNs (DIV21) by flow cytometry after staining with propidium iodide (PI) showing G0/G1, S and G2/M phases of the cell cycle. Significance was calculated using two-way ANOVA (Tukey’s multiple comparison test). ∗*p* ≤ 0.05, ∗∗*p* ≤ 0.01, *** *p* ≤ 0.001 versus controls, ns= not significant, n = 10, mean ± SEM. (B) Flow cytometry analysis results of NeuO+, Annexin V-APC and PI multicolor fluorescence staining showing FUS^WT^ and FUSP^525L^ cell apoptosis/necrosis after DMSO and 100 nM BI2536 treatment (DIV21), as described in Materials and Methods. The percentage of cells in each quadrant is presented as the mean ± SD (standard error of deviation) and the results are representative of at least six independent experiments. (C) Quantitative analysis of apoptotic and necrotic cells using APC Annexin V/PI staining flow cytometry of 72 hours in DMSO treated FUS^WT^ control and FUS^P525L^ mutant MNs showed that necrosis process was increased in FUS^P525L^ MNs compared with FUS^WT^ control MNs (D) The number of necrotic cell populations increased significantly in FUS^P525L^ MNs with 100 nM BI2536 treatment (PLK1 inhibitor) 72 hours in culture, whereas BI2536 had no effect on early or late apoptotic cell populations. For each sample, 50,000 events were measured. Dead cells were defined as early apoptotic (Annexin V^+^/PI^−^), late apoptotic (Annexin V^+^/PI^+^), and necrotic (Annexin V^−^/PI^+^). Given are the % numbers of stained cells after treatment. n = 6 independent experiments, mean ± SD. * *p* ≤ 0.05, ** *p* ≤ 0.01, *** *p* ≤ 0.001 in an Unpaired, two-tailed Student’s *t*-test, asterisks represent *p*-values compared to the controls. All FACS gating strategy figures can be found in supplementary figures.

Next, we investigated whether PLK1 upregulation might affect cell survival and regulation of cell death pathways like apoptosis or necrosis. For this, we have performed multi-color FACS analysis using Annexin V and propidium iodide (PI) labeling in FUS^WT^ and FUS^P525L^ living MNs, which had been labeled live using the neuronal fluorescent dye NeuroFluor^TM^ NeuO (Gating strategy shown in Supplementary Figure S10, S11). *FUS* mutant MNs did not show significantly increased MN cell death (Figure 6B). Finally, we asked whether cell survival is impaired in case of PLK1 inhibition. We first did a titration in proliferating FUS^WT^ and FUS^P525L^ NPCs and finally used 100 nM BI2536 treatment for further analysis, which showed significant block of cell cycle progression in both WT and mutant NPCs (Supplementary Figure S7A). Treating MNs for 72 hours with 100 nM PLK1 inhibitor BI2536 (or DMSO) reduced survival of MNs and significantly increased apoptotic and necrotic cell death (Figure 6C, D). This was seen in both FUS^WT^ and FUS^P525L^ mutant MNs, but changes in necrotic cell death rates were significantly increased in case the of the FUS^P525L^ mutation.

### PLK1 activity could be modulated by the upregulation of DNA damage response and plays a protective role in challenged cells

PLK1 has been shown to possess an increasing number of non-cell cycle related functions (*50, 51*). For example, PLK1 was recently reported to be involved in DNA damage repair in a poly(ADP-ribose) dependent manner (*52, 53*) and is therefore very similar to one of the roles recently associated with FUS function (*17*). We thus wanted to test if PLK1 upregulation could occur in response to the early upregulation of DNA damage levels in FUS^P525L^ MNs. First, we stressed generic cell culture cells with etoposide to induce DNA damage. Indeed, we found a dose dependent upregulation of PLK1 upon etoposide treatment in HEK cells in both fluorescence staining as well as western blot (Supplementary Figure S12). Excitingly, when investigating MNs, we observed a solid increase in PLK1 immunofluorescence (IF) signal in the FUS^WT^ MNs upon DNA damage induction, which is however not increased in FUS^P525L^ MNs (Figure 7A, B).

**Figure 7.**
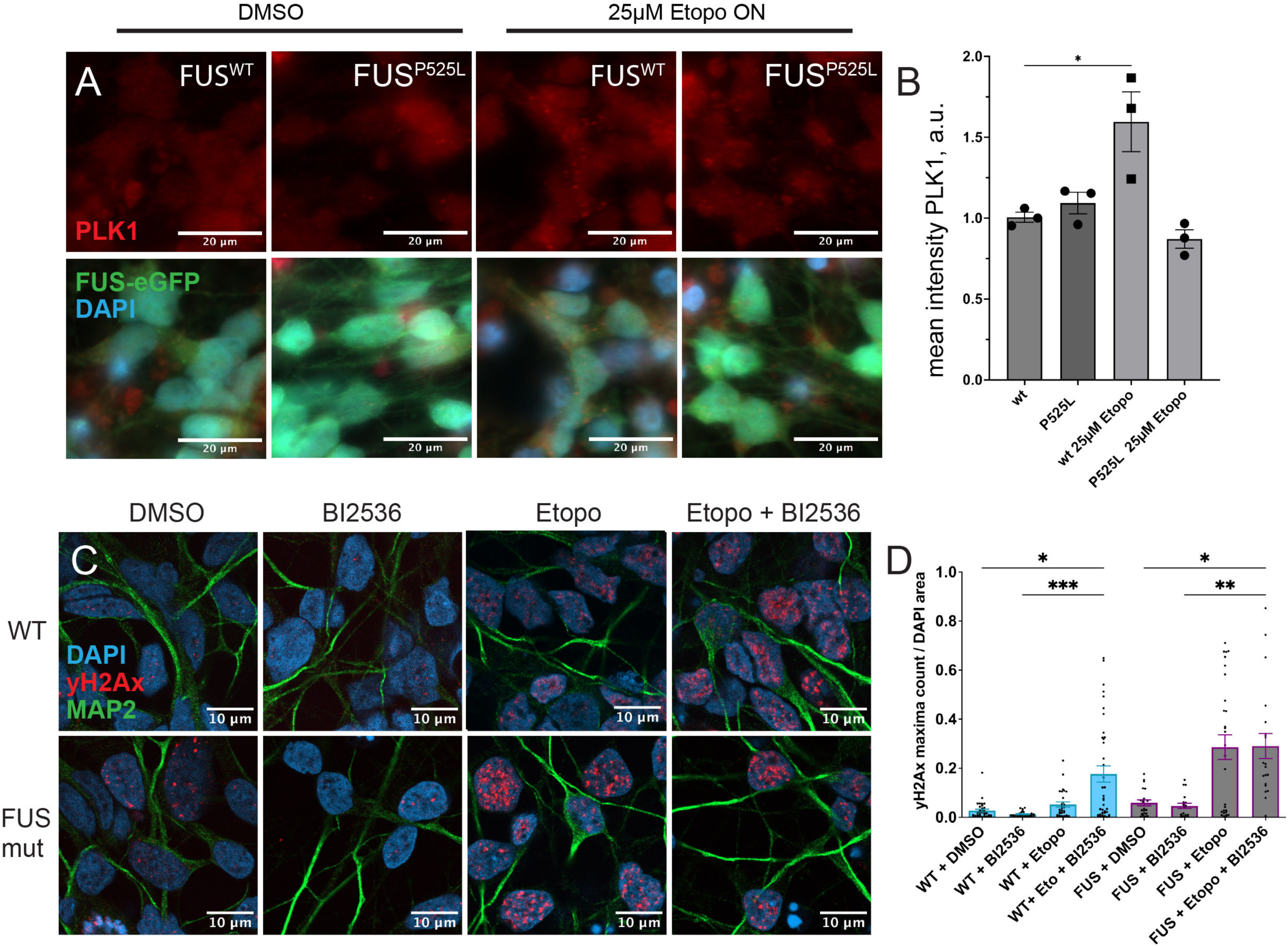
PLK1 is activated by increased DNA damage and regulates cell survival pathways. (A-B) Etoposide treatment increases overall PLK1signal in control but not in FUS^P525L^ MNs. Representative images (A) of control and FUS^P525L^ MNs treated with 25µM Etoposide and quantification (B) of mean cytoplasmic PLK1 signal per cell across all of the cells (n=3 technical replicates per condition, >3 field of view images per replica, 7-10 cells per image). (C-D) Analysis of DNA damage foci labelled by yH2AX in control and *FUS* mutant MNs, in DMSO treated cells or cell additionally treated with Etoposide or PLK1 inhibitor BI2536. Representative images (C) and box-plot of analysed yH2AX foci (D) n(exp)=3, n(cells) ≥ 19 across. Significance assessed with unequal variance *t*-test for A-B and one-way ANOVA for C-D. * *p* ≤ 0.05, ** *p* ≤ 0.01, *** *p* ≤ 0.001, **** *p* ≤ 0.0001.

We finally investigated if PLK1 activity participates in repair and coping mechanisms after DNA damage. As previously reported (*17*), we confirmed that FUS^P525L^ MNs have an increased amount of DNA damage foci labeled by γH2AX (Figure 7C, D). In unstressed conditions, inhibition of PLK1 with 1µM BI2536 for 24 hours did not affect base levels of γH2AX loci neither in control nor FUS^P525L^ MNs. When cells were further challenged with etoposide, however, both FUS^WT^ and FUS^P525L^ MNs showed a pronounced increase in DNA damage sites, with the amounts of DNA repair foci being higher in FUS^P525L^ MNs. Relevant to our hypothesis, PLK1 inhibition induced a pronounced additional increase in γH2AX loci in control cells treated with etoposide. Blocking PLK1 had, however, no significant additional effect on etoposide treatment in FUS^P525L^ MNs, possibly due to earlier activation or saturation of this pathway in these cells. Together, our data support the hypothesis of PLK1 upregulation upon DNA damage and its protective function in DNA repair and cell survival.

## Discussion

Accumulation of DNA damage and an early axonopathy with severely impaired distal axonal mitochondria is a hallmark of *FUS*-ALS. However, the reason for this axonopathy as well as the connection to DNA damage is poorly understood. We thus established a compartment-specific RNA-seq approach to identify local changes of transcriptome in axons and compared those to the SD compartments of FUS^WT^ and ALS causing FUS^P525L^ mutant MNs. Such comparison of WT and mutant *FUS*-ALS MNs identified fundamental differences between the axonal and SD compartment in general and only a little overlap between detected DEGs between SD and axonal compartments. These differences strongly support the hypothesis that axons contain very distinct sets of RNAs, which in combination with reported local axonal translation mechanisms and different metabolic environment could play an essential critical role in axonal/synapse function and can pinpoint compartment specific disease mechanisms. As an example, we detected a general increase in expression of genes that encode ECM components and cell adhesion proteins in axonal compartments in comparison to SD in both FUS^WT^ and FUS^P525L^. Several functional studies have reported that upregulation of cell adhesion proteins provides structural/functional connections between ECM molecules and intracellular components and plays a role in axonal survival, neural stem cell behavior, synaptic transmission, and cytoskeletal regulation, implying their importance in axonal plasticity and repair (*54*). Alterations in these pathways are therefore likely to be significant for ALS pathogenesis. In agreement with this, our transcriptome data showed that both SD and axonal transcripts in FUS^P525L^ MNs have a significant downregulation of pathways connected to adhesion, proteoglycan binding and ECM related processes in comparison to FUS^WT^ (Figure 3B, C, F and 4B, C, F). Thus, our results suggest a potential role of these pathways in the general axonal functions and pathogenesis of *FUS*-ALS and warrants additional investigation.

Our analysis of DEGs in SD compartment of *FUS-*ALS MNs highlighted several other pathways critical for disease development. Upregulation of DNA damage, apoptosis and mitochondrial stress pathways, as well as downregulation of neurodevelopmental processes, synaptic signaling and transmission agrees well with known early *FUS-*ALS phenotypes and cellular defects (*17, 20*). Although activation of WNT signaling has been previously implicated in neurodegenerative diseases like Alzheimer’s and Parkinson’s disease (*55*), only recently it has been also reported in various model systems of ALS and was proposed to modulate oxidative stress, mitochondrial dysfunction, autophagy, and apoptosis (*56*). Therefore, our RNA-seq data, provides an important list of candidate genes that may be relevant to better understand the role of WNT signaling in ALS and its potential therapeutic implications.

Surprisingly, axonal-specific transcriptome analysis detected only a small but well-connected set of enriched DEGs and pathways strongly focused on downregulation of proteoglycan binding and ECM related processes and upregulation of mitochondrial transcription, innate immune response signaling and PLK1/cell cycle/DNA damage repair pathways. The latter included the cell cycle and PLK1 pathway related genes including MT-associated and kinetochore proteins. Interestingly, PLK1 was recently also reported to be present in *SOD1*-, *TDP43-* and *PFN*-ALS iPSC-derived MNs, but not sensory neurons (*57*) and is a prominent marker present in many aggressive forms of cancer and some neurodegenerative diseases (*48*). PLK1 is a Ser/Thr kinase and is most known for its role in cell cycle regulation and its role in centrosome maturation, progression through mitosis and the onset of cytokinesis (*48*). However, additional roles of PLK1 include involvement in DNA replication, DNA damage response and genome stability, ER-mitochondria contact sites as well as regulating autophagy and apoptosis and inflammatory signaling (*58–60*). Importantly, most of these reports investigated PLK1’s function in cycling cells. Nevertheless, such diverse roles of PLK1 places it as a critical hub regulating many cellular processes and cell survival. Its aberrant activation therefore could be a reliable marker of cell stress and a potential therapeutic target (*61*). Comparison of our axonal datasets with other published axonal specific RNA-seq from human and mouse ALS models (*29, 35*) suggests that PLK1/cell cycle-related pathways are indeed a commonly activated pathway in axons. However, it remains unclear why MNs have this baseline expression of PLK1 even in control non-mitotic cells and what is triggering its upregulation in disease.

Based on our experiments we propose that increase in DNA damage levels in *FUS*-ALS MNs could be a potential trigger for PLK1 activation. Both HEK and control MNs are able to upregulate PLK1 levels in response to DNA damage stress. We also showed that PLK1 expression reduced the number of DNA damage foci present in healthy MNs after they were challenged with DNA damage inducing drug Etoposide. Interestingly, PLK1 is recruited to the double strand breaks induced by laser ablations and such recruitment depends on poly (ADP-) ribose polymerase 1 (PARP-1) and is slowly reversed in PARG-dependent manner (*52*). Protective role of PLK1 in the MNs could be connected to PLK1 activation in response to DNA damage. Interestingly very similar recruitment dynamics and PARP-1/PARG regulation has been also demonstrated for FUS and could potentially explain PLK1 activation in the absence of nuclear FUS (*17, 62*). Whether PARP1-dependent PLK1 recruitment to DNA damage sites could potentially compensate for the lack of FUS recruitment or if these mechanisms are independent remains to be tested. Furthermore, the compartment specific upregulation of PLK1 needs also further investigation. Relevantly, a very recent study reported that FUS is involved in mitochondrial DNA repair and *FUS*-ALS mutations disturb mitochondrial DNA damage repair (*22*). Of note, mitochondrial damage is mainly found in distal axons in case of *FUS*-ALS. Thus, it will be important to determine whether PLK1 is also involved in mtDNA damage repair associated pathways.

Activation of PLK1 can be connected to the coping mechanism of the affected cells. Inhibition of PLK1 in FUS^P525L^ led to significantly elevated MNs death and diminished their already reduced viability and survival. This is reminiscent to outcomes in many cancer cell model systems, where PLK1 inhibition compromises survival of cancer cells through various mechanisms including inducing proteasome activity and reducing ubiquitination of proteins (*63, 64*). Notably, disturbance of protein homeostasis is a key hallmark and perhaps determinant of numerous neurodegenerative disorders, including ALS (*13, 65*). Our data does not specifically determine if PLK1 has a direct role in downregulating these cellular processes, and whether specific cell death pathways were involved. However, recent evidence strongly implicated ferroptotic cell death upregulation in several ALS models including *FUS*-ALS, which is a form of necrotic cell death (*66, 67*). Along this line, PLK1 function has been shown to inhibit ferroptosis in esophageal squamous cell carcinoma through activity of pentose phosphate pathway and regulation of NADPH and glutathione levels (*63*).

Alternative hypothesis for downstream mechanisms would involve potential role of PLK1 in regulating cellular innate immune response mechanisms in ALS (*68*). PLK1 activity has been shown to inhibit interferon activation via phosphorylation and inhibition of the mitochondrial protein MAVS (*69*). Furthermore, inhibition of either PLK1 or HDAC, has been shown to activate interferon immune response reporter RIG-I in cell reporter assay (*70*). The latter results are especially intriguing, since HDAC1 has been shown to bind FUS and recruit it to double-stranded DNA-break repairs, while inhibition of HDAC1 has protective roles in ALS models (*71, 72*). Our observation of PLK1 upregulation therefore could be an important link connecting different DNA damage response mechanisms among each other and/or to potentially activate innate immune response in (FUS-) ALS MNs. Such hypothesis is supported by the fact that one of the few identified PPI networks in our distal axonal analysis maps to cluster connected to the immune response reporter RIG-I/interferon/inflammation and Herpes simplex virus 1 infection pathways. The latter would also fit the hypothesis of retrotransposon and virus reactivation as potential contributing mechanisms of ALS (*28, 73*). In addition to the immune-related enriched terms from the PPI analysis, we also identified a network related to activation of mitochondrial transcription/metabolism pathways in FUS^P525L^ mutant motor axons, as compared to the healthy controls, and GSEA further showed increased expression across the mitochondrial functional gene set. This finding was intriguing given that prior studies in MNs and postmortem tissues have demonstrated crucial function of FUS in maintaining mitochondrial DNA/RNA integrity following overexpression of FUS and through its involvement in repair mechanisms (*17, 18, 22, 74, 75*). Therefore, additional studies are required to address protective mechanisms underlying mitochondrial function and transcriptional fidelity in FUS^P525L^ MN axons, which are highly reliant on mitochondria to meet their energy demands. Many of the genes from the PLK1 associated cluster, like KIF2C, BUB1B, CENPM, CENPE, and PARPBP, have been increasingly implicated in axon growth and regeneration via regulating axon morphology, (*76, 77*). Abnormal upregulation of these genes could be behind reported abnormalities in cytoskeleton morphology and axonal transport defects and could be used as a potential target for disease mitigation strategy. As an example, both pharmacological inhibition and shRNA knockdown of AuroraB was shown to enhance mitochondrial transport in iPSC derived MNs from SOD1^A4V^ ALS patients (*78*). PLK1 inhibition was able to rescue neurofilament morphology and axonal phenotypes in Charcot Marie Tooth type E iPSC-derived neurons (*79*). Furthermore, Protein Phosphatase 2A (PP2A) is a phosphatase required for the PLK1 inactivation during DNA damage response (*80*). Its inhibition or downregulation and therefore supposed activation of PLK1 was found in modifier screens to rescue mitochondrial trafficking defects and neuromuscular junction failure in *FUS* mutant flies and *FUS*-ALS iPSC-derived MNs (*81*). All above mechanisms could potentially provide a link between DNA damage response and regulation of cytoskeleton in healthy MNs, as well as understanding how aberrant expression of these genes contributes to axon’s vulnerability in disease.

In summary, our results highlight the alteration of many pathways in our model FUS^P525L^ MNs, confirming previously reported critical pathways, such as DNA damage and cellular metabolism, as well as identifying a number of new potential target genes that could be relevant in our understanding and treatment of the disease. We propose that PLK1 pathway activation is a response to the increased DNA damage observed in these cells and could provide a protective mechanism for the survival of these affected cells (Figure 7). It will be essential to further investigate what are the consequences of these pathway activation and if it can be modulated to improve survival and health of the affected cells. In addition, PLK1 activation could provide a potential link between increased DNA damage, activation of inflammatory pathways, distal mitochondrial axonal phenotypes and compromised cell survival in *FUS*-ALS MNs, which deserves further systematic functional studies. It becomes evident that some prominent underlying symptoms and biomarkers are present in patients from early on and our focus on misregulated pathways we present here would be critical for our potential ability to detect, modulate and prevent the disease.

## Supplementary Figures

**Supplemental Figure S1. Analysis of SD and axon FUS^P525L^ and FUS^WT^ MNs on normalized expression values of expressed genes.**

(A) PCAs of gene expression (top 500 most diverse genes) reveals clear separation of SD and axonal datasets.

(B and C) Heatmaps showing the pair-wise (B) Pearson (distance based) (C) Spearman’s rank (ranking based) correlation between all soma samples.

(D and E) Heatmaps showing the pair-wise (D) Pearson (distance between genes) and (E) Spearman’s rank (ranking of the genes) correlation between all axon samples.

(F and G) Heatmap of genes with the highest mean expression strength across (F) soma samples and (G) axon samples. The scale goes from blue to red representing the highest expression.

**Supplementary Figure S2. Functional enrichment analysis between SD and axonal transcriptomes highlights axonal specific pathways.**

(A-C) Gene ontology (GO; biological process (BP), molecular function (MF), and cellular component (CC)) in (A) upregulated, (B) downregulated, and (C) pathway analysis of DEGs in FUS^WT^ control MNs. (D-F) GO Functional analysis on (D) upregulated, (E) downregulated, and (F) pathway analysis of DEGs in FUS^P525L^ mutant MNs. GO/pathway enrichment analysis were analyzed by Metascape and EnrichR, respectively. Significant terms are listed with a *p*-value ≤ 0.05 significance. Three main GO categories and pathways were shown in different color bars (orange: Biological Process, green: Molecular Function, violet: Cellular Component, Blue/Pink: Pathways). (see Supplementary Table S3).

(G) Cross-comparison of the axon-specific DEGs from the current study with the human axon-specific DEGs from control samples of iPSC-derived MNs (Nijssen, Aguila et al. 2018)

(H) and the primary mouse MN axons and mESC-derived MN axonal datasets from control cells (*29, 35*) revealed that a vast majority of the transcripts identified with axonal DEGs overlapped with the other datasets (an overlap of 453 and 60 genes). Venn diagram depicting the overlap revealed that these genes enriched in pathway terms across datasets included RNA metabolism, local translation, PLK1 pathway and neuronal growth/survival functions. Pathways with a *p*-value of ≤ 0.05 were considered statistically significant. (see Supplementary Table S5).

**Supplemental Figure S3. Gene set enrichment analysis (GSEA) against the GO and KEGG database for differentially enriched terms and pathways between axon versus SD of FUS^WT^ MNs.** (A-L) GSEA plots were generated using normalized counts from the DESeq2 output ranked list in axon versus SD, with a adjusted *p*-value ≤ 0.05 cutoff. Based on normalized enrichment score (NES) values, the top enriched GO terms and pathways in the FUS^WT^ group were shown. NES value represents the enrichment score after normalization. The higher the NES value, the more genes were enriched in the pathway. The y-axis represents enrichment score for the overall gene set and on the x-axis are genes (vertical black bars) represented in gene sets, indicating where the members of the gene set appear in the ranked list of genes. GSEA analysis was performed using *Partek*^TM^ *Flow*^TM^ software, v11.0.

**Supplemental Figure S4. Gene set enrichment analysis (GSEA) against the GO and KEGG database for differentially enriched terms and pathways between axon versus SD of FUS^P525L^ MNs.** (A-H) GSEA plots were generated using normalized counts from the DESeq2 output ranked list in axon versus SD, with a adjusted *p*-value ≤ 0.05 cutoff. Based on NES values, the top enriched GO terms and pathways in the FUS^P525L^ group were shown. NES value represents the enrichment score after normalization. The higher the NES value, the more genes were enriched in the pathway. The y-axis represents enrichment score for the overall gene set and on the x-axis are genes (vertical black bars) represented in gene sets, indicating where the members of the gene set appear in the ranked list of genes. GSEA analysis was performed using *Partek^TM^ Flow^TM^* software, v11.0.

**Supplemental Figure S5. Gene set enrichment analysis (GSEA) against the GO and KEGG database for differentially enriched terms and pathways in the SD of FUS^P525L^ and FUS^WT^ MNs.** (A-H) GSEA plots were generated using normalized counts from the DESeq2 output ranked list in SD, with a adjusted *p*-value ≤ 0.05 cutoff. GSEA identified GO terms and pathways related to (A and B) negative regulation of nervous system/neuron development, (C-E) mitochondrial apoptotic process, (F) trans-Golgi network, and (G and H) proteolysis process and deubiquitinase activity. The y-axis represents enrichment score for the overall gene set and on the x-axis are genes (vertical black bars) represented in gene sets, indicating where the members of the gene set appear in the ranked list of genes. GSEA analysis was performed using *Partek*^TM^ *Flow*^TM^ software, v11.0.

**Supplemental Figure S6. Gene set enrichment analysis (GSEA) against the GO and KEGG database for differentially enriched terms and pathways in the axon of FUS^P525L^ and FUS^WT^ MNs.** (A-J) GSEA plots were generated using normalized counts from the DESeq2 output ranked list in axon, significance threshold set at a adjusted *p-*value ≤ 0.05. GSEA identified GO terms and pathways related to (A and B) DNA repair pathway, (C-G) mitochondrial gene expression/metabolism functions, (H) focal adhesion, and (I and J) MAPK and PI3K-Akt signaling pathways. The y-axis represents enrichment score for the overall gene set and on the x-axis are genes (vertical black bars) represented in gene sets.

**Supplemental Figure S7. Inhibition of PLK1 with BI2536, a potent and selective inhibitor, disrupts cell proliferation and induces mitotic defects in NPCs but does not trigger MN differentiation.** (A) Representative histograms of propidium iodide (PI) fluorescence showing distribution of cell cycle phases in the DMSO and BI2536 treated samples between FUS^WT^ and FUS^P525L^ NPCs; FACS profile of exponentially growing NPC cells treated for 24 hours by BI2536 with concentrations ranging 3 nM-1 μM. After 24 hours, the NPCs were harvested and the DNA content was analyzed with flow cytometry. Inhibition of PLK1 (BI2536-treated NPCs) by flow cytometry confirm a clear accumulation of 4*N* DNA content, promoting G2/M arrest in a dose-dependent manner. The positions of 2N and 4N DNA content are indicated. Mitotic arrest is evident already at concentrations of 100 nM, thus indicating a complete mitotic arrest followed by cell death. (B) Representative images of FACS gating strategy demonstrating flow cytometric analysis of DNA cell cycle after propidium iodide (PI) staining in NPCs.

**Supplemental Figure S8. Flow cytometric gating analysis of DNA cell cycle after PI staining in FUS^WT^ MNs.** Representative images of FACS gating strategy demonstrating flow cytometric analysis of DNA cell cycle after propidium iodide (PI) staining in MAP2 positive FUS^WT^ MNs (fixed with 70% ethanol) under untreated conditions. In the illustration, the FlowJo software enables a clear classification/quantification into the cell cycle phases (G0/G1, S, G2/M) based on the PI content of the cells. Cell cycle histograms were performed by selecting single-cell events with unfractionated DNA only.

**Supplemental Figure S9. Flow cytometric gating analysis of DNA cell cycle after PI staining in FUS^P525L^ MNs.** Representative images of FACS gating strategy demonstrating flow cytometric analysis of DNA cell cycle after propidium iodide (PI) staining in MAP2 positive FUS^P525L^ MNs (fixed with 70% ethanol) under untreated conditions. In the illustration, the FlowJo software enables a clear classification/quantification into the cell cycle phases (G0/G1, S, G2/M) based on the PI content of the cells. Cell cycle histograms were performed by selecting single-cell events with unfractionated DNA only.

**Supplemental Figure S10. Flow cytometry apoptosis/necrosis multicolor staining analysis in FUS^WT^ MNs.** (A) Flow cytometry apoptosis/necrosis multicolor staining results of (A) DMSO control (B) BI2536 (100 nM), (C) unstained, (D) NeuO+ groups in FUS^WT^ MNs. Representative pseudocolor plots in MNs illustrating quantitative analysis of cell death using flow cytometric APC Annexin V/PI staining assay. Briefly, FUS^WT^ and FUS^P525L^ MNs were incubated with buffer or the indicated concentration of BI2536 (100 nM). After 72 hours, the MNs were labeled with NeuroFluor™ NeuO dye (1:1000) and incubated for 1 hour in culture media at 37 °C. After 1 hour, the MNs were collected and cell death (APC Annexin V/PI) was analyzed with flow cytometry.

**Supplemental Figure S11. Flow cytometry apoptosis/necrosis multicolor staining analysis in FUS^P525L^ MNs.** (A) Flow cytometry apoptosis/necrosis multicolor staining results of (A) DMSO control (B) BI2536 (100 nM), (C) unstained, (D) NeuO+ groups in FUS^P525L^ MNs. Representative pseudocolor plots in MNs illustrating quantitative analysis of cell death using flow cytometric APC Annexin V/PI staining assay. Briefly, FUS^WT^ and FUS^P525L^ MNs were incubated with buffer or the indicated concentration of BI2536 (100 nM). After 72 hours, the MNs were labeled with NeuroFluor™ NeuO dye (1:1000) and incubated for 1 hour in culture media at 37 °C. After 1 hour, the MNs were collected and cell death (APC Annexin V/PI) was analyzed with flow cytometry.

**Supplemental Figure S12. Etoposide treatment upregulates PLK1 in HEK cells.** (A-B). Representative images (A) and relative intensity quantification (B) of immunostaining for PLK1 protein in HEK control cells or treated with increasing doses of DNA damage-inducing drug Etoposide. (C-D). Representative western blot images (C) and its quantifications (D) showing upregulation of total PLK1 protein but no changes in phosphorylation of conserved tyrosine (T210) in PLK1 protein after Etoposide treatment. Significance assessed with unequal variance *t*-test. * *p* ≤ 0.05, ** *p* ≤ 0.01, *** *p* ≤ 0.001, **** *p* ≤ 0.0001.

## Materials and methods

**Supplementary Table S10:** Cell lines used in this study.

| Genotype | Cell line | Sex | Age at biopsy | Mutation | Primarily characterized in |
| --- | --- | --- | --- | --- | --- |
| Isogenic pair control1 | FUS<sup>WT</sup>-eGFP<sup>het</sup> | isogenic to FUS<sup>R521C</sup> and FUS<sup>P525L</sup> eGFP | N/A | - | (17) |
| Isogenic pair Mut1 | FUS<sup>P525L</sup>-eGFP<sup>het</sup><br>FUS<sup>R521C</sup> het | isogenic to FUS<sup>R521C</sup> and FUS<sup>WT</sup> eGFP female | N/A | P525L | (17, 38) |
| Control 2 | 2062-2 | male | 43 |  | (82) |
| Control 3 | GM23251-4 | female | 41 |  | (83) |
| Control 4 | AKC26 | female | 48 |  | (84) |
| Mut2 | FUS<sup>R521C</sup> | female | 58 | R521C | (17, 38) |
| Mut3 | R495QfsX527 | male | 46 |  | (17, 38) |

### Cell lines

For RNA-seq we used the two pairs of isogenic cell lines FUS^WT^-eGFP and FUS^P525L^-eGFP that were generated, described and characterized in Naumann, Pal, *et al*., 2018 and Marrone *et al*., 2018. MNs were generated following the protocol established and used in these papers.

All other cell lines were also from our previous studies and were obtained from skin biopsies of patients or healthy volunteers or generated using CRISPR/Cas9 mutagenesis and have been described before (Supplementary Table 10). The performed procedures were in accordance with the Declaration of Helsinki (WMA, 1964) and approved by the Ethical Committee of the Technische Universität Dresden, Germany (EK 393122012 and EK 45022009) and by the Swedish Ethical Committee for Medical Research (Nr 94-135 to perform genetic research, including for *FUS*, and Nr 2018-494-32M to perform in vitro lab studies on cell lines derived from patients with ALS). Written informed consent was obtained from all participants, including for publication of any research results. All fibroblast lines were checked for Mycoplasma spp. before and after reprogramming, and afterward, routine checks for Mycoplasma were conducted every three to six months. We used the Mycoplasma detection kit according to the manufacturer’s instructions (Firma Venor GeM, No 11–1025).

### Differentiation of NPCs to MNs

The generation of human NPCs and MNs was accomplished following the modifications protocol from Reinhardt *et al.* (2013) and described also in Naumann, Pal, *et a*l., 2018 (*17, 37*). In brief, iPSCs colonies were collected and stem cell medium, containing 10 µM SB-431542, 1 µM Dorsomorphin, 3 µM CHIR 99021 and 0.5 µM SAG (Cayman; 11914) was added. After 2 days hESC medium was replaced with N2B27 consisting of the aforementioned factors and DMEM-F12/Neurobasal 50:50 with 1:200 N2 Supplement, 1:100 B27 lacking Vitamin A and 1% penicillin/streptomycin/glutamine. On day 4 150 µM ascorbic acid was added while Dorsomorphin and SB-431542 were withdrawn. 2 days later the EBs were mechanically separated and replated on Matrigel coated dishes. For this purpose, Matrigel was diluted (1:100) in DMEM-F12 and kept on the dishes over night at room temperature. Possessing a ventralized and caudalized character the arising so called small molecule NPCs (smNPC) formed homogenous colonies during the course of further cultivation. It was necessary to split them at a ratio of 1:10–1:20 once a week using Accutase for 10 minutes at 37 °C.

Final MN differentiation was induced by treatment with differentiation medium with 0.5 µM SAG, 0.2mM Ascorbic Acid, 1 µM retinoic acid (RA), 1ng/ml BDNF, 1ng/ml GDNF 0.5 µM SAG in N2B27 and changed every 2 days. After 6 days differentiation media was changed to maturation media containing 5ng/ml Activin A, dbcAMP 0.1mM, 2ng/ml BDNF, 2ng/ml GDNF, 1ng/ml TGFß-3, 0.2mM Ascorbic Acid. On day 7-10 another split step was performed to seed them on a desired cell culture system and kept in the maturation media without Activin A but with one-time addition of 2.5 µM DAPT. Following this protocol, it was possible to keep the cells in culture for over 2 months.

### Growth of MNs in microfluidic chambers

The MFCs were purchased from Xona (RD900 with 900 μm long microgrooves). At first, Nunc glass bottom dishes with an inner diameter of 27 mm were coated with Poly-L-Ornithine (Sigma-Aldrich P4957, 0.01% stock diluted 1:3 in PBS) overnight at 37 °C. After 3 steps of washing with sterile water, they were kept under the sterile hood for air drying. MFCs were sterilized with 70% Ethanol and also left drying. Next, the MFCs were dropped onto the dishes and carefully pressed on the glass surface for firm adherence. The system was then perfused with Laminin (Roche 11243217001, 0.5 mg/ml stock diluted 1:50 in PBS) for 3 hours at 37 °C. For seeding cells, the system was once washed with medium and then 10 μl containing a high concentration 3 ×10^5^ cells (3 × 10^7^ cells/ml) were directly injected into the main channel connecting two wells. After allowing for cell attachment over 30–60 minutes in the incubator, the still empty wells were filled up with maturation medium. This method had the advantage of increasing the density of neurons in direct juxta-position to microchannel entries whereas the wells remained cell-free, thereby reducing the medium turnover to a minimum. To avoid drying out, PBS was added around the MFCs. Two days after seeding, the medium was replaced in a manner which gave the neurons a guidance cue for growing through the microchannels. Specifically, a growth factor gradient was established by adding 100 μl N2B27 with 500 μM dbcAMP to the proximal seeding site and 200 μl N2B27 with 500 μM dbcAMP, 10 ng/μl BNDF, 10 ng/μl GDNF and 100 ng/μl NGF to the distal exit site. The medium was replaced in this manner every third day. After 7 days, the first axons began spreading out at the exit site and cells were typically maintained for up to six weeks.

### RNA extraction and sequencing analysis

For transcriptome sequencing (RNA-seq), MNs of the cell lines FUS^WT^ and FUS^P525L^ were generated in biological triplicates (3 independent differentiations side by side) and two-week-old microfluidic chambers (MFC) with abundant outgrowth of axons were selected for RNA isolation. A total of 7-10 MFC were used per each cell line condition to extract the total RNA. Briefly, both SD and axonal compartments were washed with PBS and were lysed by direct application of Qiazol lysis buffer (Qiagen, Hilden, Germany) from the kit into chambers. Following that, RNA purification was performed using an RNeasy Mini Kit (Qiagen, Hilden, Germany) according to the manufacturer’s instructions including a column DNase digest for all samples. RNA quantity and quality was evaluated by Agilent 2100 Bioanalyzer using Agilent RNA 6000 Pico Kit (Agilent Technologies, CA, USA). For RNA quality assessment, fragment size distribution (DV200; percentage of RNA fragments with a length > 200 nucleotides) was calculated for each sample using 2100 Expert software. Samples with DV200 ≥ 60% were subjected to RNA seq. mRNA was purified from 10 ng total RNA and exponentially amplified using Takara Bio SMARTer-Seq (Clontech/Takara Bio USA) according to the manufacturer’s instructions. RNA libraries for both SDs and axon RNA-seq were prepared using a SMARTer HV chemistry, Ultra mRNA-Seq Library Prep Kit-v3 (Clontech/Takara Bio USA) following manufacturer’s recommendations and processed on an Illumina HiSeq 2500 system (polyA[+] RNA-seq), resulting in 75 bp single-end sequencing mode to an average depth of 30 million fragments per library. Sequencing libraries were constructed and sequenced at next-generation sequencing facility (Dresden-concept Genome Center), on Illumina HiSeq2500 platform.

FastQC (v0.11.3) measure was used to perform a basic quality control (QC) on the sequence data. Reads were mapped to the human reference genome hg38/GRCh38 (obtained from EnsEMBL v81) using the aligner GSNAP (v2017-11-15) (with splice-junction support from annotated genes (EnsEMBL v81)) (*85*). Further quality control on mapped reads, rRNA content, coverage of exonic/intronic/intergenic regions and number of detectable genes was done with RNA-SeQC (v1.1.8) (*86*). A table of raw read counts per gene was obtained based on the overlap of the uniquely mapped reads with annotated human genes (EnsEMBL v81) using featureCounts (v1.5.2) (*87*). Normalization of the raw read counts based on library size was performed with the DESeq2 R package (v1.16.1) (*88*). Principal component analysis (PCA), sample-to-sample Euclidean distance as well as Pearson’s and Spearman’s correlation coefficients were computed based on the normalized gene expression level using R. For testing for differential expression with DESeq2, the count data was fitted to the negative binomial distribution and adjusted *p*-values for the statistical significance of the fold change (FC) were adjusted for multiple testing with the Benjamini-Hochberg correction for controlling the false discovery rate (FDR) accepting a maximum of 5% false discoveries (adjusted *p*-value ≤ 0.05) and log2fold change (FC) ≥ 1. Heatmap and unsupervised hierarchical clustering of DEGs in axon and soma samples of control/mutant MNs were generated using R package. Heatmap and unsupervised hierarchical clustering of DEGs in axon and soma samples of control/mutant MNs were generated using R and *Partek*™ *Flow*™ software, v11.0. Data are deposited at NCBI/GEO under the accession number GSE276214.

### Use of published datasets

All data used in cross-comparisons of datasets were obtained from the NCBI/GEO database. The GEO accession numbers and their respective study references for the axonal transcriptomic studies were as follows: GEO: GSE66230 (*29*), GEO: GSE121069 (*35*) (Supplementary Table S5). In all cases for RNA-seq, FASTQ data was processed and DESeq2 normalization values were calculated using the pipeline described above in *Partek*^TM^ *Flow*^TM^ software, v11.0, to avoid differential mapping bias across sample datasets.

### Functional annotation of DEGs by GO and KEGG analysis

To better interpret the possible pathways and target genes of the relevant profiles of the SD and axon-enriched genes analysed in *FUS*-ALS, GO functions and pathway (Reactome and Kyoto Encyclopedia of Genes and Genomes (KEGG)) enrichment analyses of candidate genes were carried out by utilizing the Metascape and EnrichR, integrated online tools for gene list annotation and biological analysis (*89, 90*). Metascape integrates several functional databases, such as GO, KEGG and Uniprot to analyze multiple gene sets simultaneously. The DEGs were analyzed and visualized by Metascape with the criteria of minimum overlap ≥ 3, *p*-value ≤ 0.05, and minimum enrichment score > 1.5. We utilized EnrichR (enrichment testing across all GO functions, pathways) with Fisher’s exact test to conduct the functional enrichment analysis with the DEGs. In Metascape and EnrichR analysis, we combined the signaling pathways from two libraries, including KEGG and Reactome, to create a single analysis route. Only the important paths for which the *p*-values ≤ 0.05 were evaluated and considered after deleting duplicate pathways. For functional GO annotations, we looked at the GO biological process (BP), GO molecular function (MF), and GO cellular component (CC) datasets in Metascape and EnrichR and selected the most important GO terms based on set criteria and with a *p*-value ≤ 0.05. Gene set enrichment analysis (GSEA) (*41*)was performed on the normalized gene counts (DESeq2) of RNA-seq data (mutant and control) using the GSEA function in the *Partek*^TM^ *Flow*^TM^ software, v11.0. The used gene sets for testing were the GO-term derived gene set database (BP, MF, and CC) and KEGG pathway database, and the significant gene sets were defined as those with a normalized enrichment score (NES) > 1 and *p* ≤ 0.05.

Venn diagrams of target genes showing axon-soma differential expression (FC ≥ 1 or FC ≤ −1, adjusted *p*-value ≤ 0.05) in ALS model cultures were selected as the candidate target genes for further downstream analysis and extracted via the online tool InteractiVenn.

### PPI network construction and identification of key genes

The protein-protein interaction (PPI) networks were generated with the Search Tool for the Retrieval of Interacting Genes (STRING) database v12.0 as well as stringApp of Cytoscape v3.10.1 software (*42, 91*). In the present study, PPI network of axon (137) and SD (166) differentially expressed genes (DEGs) was generated with a medium confidence score of 0.4 (confidence score ≥ 0.4), disconnected nodes were excluded from the network. To further strengthen the PPI core network of identified SD upregulated and axon downregulated DEGs, 5-10 direct binding partners were added to reveal functional interactions between the deregulated proteins. Functional enrichment analyses were carried out using STRING database. The nodes indicate the DEGs and the edges indicate the interaction (databases, high-throughput experiments, co-expression/co-occurrence, text mining, neighbourhood genes) between two proteins. The PPI network was subsequently imported into Cytoscape for further visualization and analysis. Furthermore, the rank and score of these key genes/clusters were identified based on a network topological property using NetworkAnalyzer v4.4.8 plugin of the Cytoscape. The statistical enrichment analyses in STRING indicated that the DEGs were significantly enriched in PPI networks (*p*-value ≤ 0.05).

### Cell cycle analysis

Cell cycle analysis for cultured NPCs was performed after 24 hours of incubation with DMSO or BI2536 (a potent and selective PLK1 inhibitor, Selleckchem) treatment with concentrations ranging from 3 nM-1000 nM. Briefly, the cells (∼1-2 × 10^6^ cells/ml) were collected in 500 µL and fixed by the dropwise addition of 500 µL of ice cold 70% ethanol (−20 °C) whilst gently mixing and stored at 4 °C overnight. The next day, the ethanol-PBS 1:1 solution was removed by pelleting the cells (300 × g, 5 minutes, 4 °C), followed by two washing steps with cold PBS. The cells were resuspended in PBS containing 0.5 mL/tube FxCycle™ PI/RNase staining solution (Propidium iodide/RNase, Thermo Fisher Scientific) following the manufacturer’s guidelines and incubated in the dark at 37 °C for 30 minutes. For each sample 50,000 events were acquired and determination of NPCs distribution among G0/G1, S, G2/M cell cycle phases of each group was performed at 488 nm using a FACSCalibur flow cytometer (BD Biosciences, San Jose, CA, USA). Data were analyzed by BD FlowJo software v10.7.2 (BD Pharmingen, San Diego, CA, USA) and the gating strategy is shown in Supplementary Figure S7.

For cultured MNs (DIV21), the cells (∼1-2 × 10^6^ cells/sample) were harvested (Accutase (Sigma) for 5 min at 37 °C) and washed twice with PBS. All procedures were performed using FACS buffer (PBS, 1 mM EDTA, 5% normal goat serum, 10% FBS). After addition of FACS buffer, the cell suspension was centrifuged (300 × g, 5 minutes, 4 °C), washed twice with cold PBS, and fixed in ice-cold 70% ethanol. Intracellular staining was done after 2 hours of fixation and permeabilization with ethanol at 4 °C using the polyclonal chicken anti-MAP2 primary antibody (1:2000, Abcam) and incubated (in 100 µl FACS buffer) for 60 minutes on ice. The cells were washed with FACS buffer and labeled with a secondary Alexa Fluor 647-conjugated (infrared) antibody (goat anti-chicken IgY, 1:1000, Invitrogen) and incubated again for 45 minutes on ice. In order to exclude non-specific binding of the Alexa Fluor-labeled secondary antibody, control cells were additionally stained with the secondary antibody without using the primary antibody. For MAP2 quantification, the number of cells that were positive for the secondary antibody was subtracted from the MAP2+ secondary antibody stained cells. Cells were washed and resuspended in 500 µl FACS staining/buffer solution consisting of 500 µl/sample FxCycle™ PI/RNase staining solution (Thermo Fisher Scientific) according to the manufacturer’s instructions and incubated in the dark at 37 °C for 30 minutes. Finally, flow cytometry measurements were taken at excitation wavelength of 488 nm using a FACSCalibur flow cytometer (BD Biosciences, San Jose, CA, USA). For each sample 50,000 events were acquired and data analysis to detect the MAP2-positive MNs cell cycle (G0/G1, S, G2/M) distribution of each group was performed using BD FlowJo v10.7.2 software (BD Pharmingen, San Diego, CA, USA). Specific subpopulations as well as related flow cytometry gating scheme were defined as outlined in the supplementary Figures S8, S9.

### Apoptosis/necrosis analysis

Apoptotic/necrotic cells were quantified by flow cytometry using an Annexin V/PI and NeuO multicolor staining procedure (BioLegend and STEMCELL Technologies) according to the manufacturer’s instructions. In brief, DMSO or BI2536 (100 nM, Selleckchem, 72 hours, 37 °C) treated MNs (DIV21) were labelled with NeuroFluor™ NeuO dye (STEMCELL Technologies) with Ex/Em of 468/557 nm that selectively labels neurons in live culture. The dye was diluted in the respective culture medium to the concentration of 0.1 µM and the medium was applied directly to the cells, which were then incubated for 1 hour at 37°C. Next, the labeling medium was removed and replaced with fresh culture medium and the cells were incubated for another 90 minutes. After that, the cells were washed once with FACS buffer (PBS, 1 mM EDTA, 5% normal goat serum, 10% FBS) and detached with Accutase (Sigma, 5 minutes at 37 °C). The detached cells (from culture medium and Accutase treatment) were then washed twice with cold FACS buffer (300 × g, 5 minutes, 4 °C) and the cell pellet was resuspended at the density of 1 × 10^6^ cells/ml in 100 µl Annexin V binding buffer (BioLegend, #422201). All the following steps were performed on ice. Cell suspension was then filtered through the cell strained cap with 35-µm nylon mesh. Subsequently, 5 μl APC Annexin V (BioLegend, #640919) was added to cell suspension and the cells were incubated for 15 minutes in the dark (RT), followed by the addition of 10 µl of 0.5 mg/ml PI (BioLegend, #421301) to identify dead cells and incubation of the cells for 5 minutes at 4 °C. After incubation period, 400 μl Annexin V binding buffer was added and APC Annexin V fluorescence and PI fluorescence were then detected using a Cytek^TM^ Aurora spectral flow cytometer running on the SpectroFlo software v2.2.0.3 (Cytek Biosciences, Fremont, CA, USA). At least 50,000 events were recorded for each sample and the acquired apoptosis/necrosis data were analysed using FlowJo v10.7.2 software (BD Pharmingen, San Diego, CA, USA). Gating strategies are shown in Supplementary Figures S10, S11. Unstained and single stained cells were used as controls and measured in every single experiment. Annexin V^−^/PI^−^ cell populations were indicated living cells, Annexin V^+^/PI^−^ were considered early apoptotic cells, whereas those that were positive for both Annexin V^+^/PI^+^ were identified as necrotic or late apoptotic cells. The apoptotic rates were determined as the percentage of the total cell population. All FACS measurements including cell cycle analysis were carried out in the Core Facility for Cell Sorting and Cell Analysis, University Medical Centre Rostock, Germany.

### qPCR PLK1 mRNA

mRNA was prepared from a DIV21 cultures of control and P525L MN cultures grown in microfluidic chambers using Qiagen mRNA mini kit. cDNA was generated using Superscript RT DNA polymerase (Thermo Fischer Scientific). We used IQ SYBR Green Supermix (Biorad) and mRNA preps from 3 separate mRNA samples were run in triplicates, and were normalized against two different human house-keeping genes (RBS18 and HPRT). Following primers were used: PLK1: GCACAGTGTCAATGCCTCCAAG and GCCGTACTTGTCCGAATAGTCC, RBS18: GCAGAATCCACGCCAGTACAAG and GCTTGTTGTCCAGACCATTGGC and HPRT: CATTATGCTGAGGATTTGGAAAGG and CTTGAGCACACAGAGGGCTACA.

### Western blot

Protein samples for Western blot were prepared from MN cultures grown in multi well plates and lysed using Mild protein lysis buffer and sonication. Standard mild lysis buffer is 50 mm Tris-HCl, pH 7.8, 150 mm NaCl, 1% Triton X-100, 10% glycerol, 25 mm NaF, 1× Roche cOmplete^TM^ Mini tablets protease inhibitor (5-10 µM is usually sufficient). Samples were prepared from independent experiments. Purified anti-PLK1 Antibody (Mouse) (1:1000) (#627701, Biolegend) and PLK1 (208G4) Rabbit mAb #4513 (Cell Signaling technology, 4513S) (1:500). Either Mouse a-Tubulin antibody or GAPDH (14C10) Rabbit mAb (HRP Conjugate) #3683 Cell Signaling was used as a loading controls. Secondary antibodies were either IRDye® 800CW Goat anti-Rabbit IgG Secondary Antibody and IRDye®, 680RD Goat anti-Mouse IgG Secondary Antibody (LicorBio) or HRP-conjugated anti-mouse and anti-rabbit with ECL detection reagents.

### Cell culture and treatments

Etoposide treatment of HEK cells and MNs: HEK cells were grown in DMEM/F12, 10% FBS, Pen/Strep media with Non-essential amino acids (from 100x stock) and Sodium Pyruvate (100x stock). Cells were treated with 5 µM, 25 µM or 50 µM Etoposide and either lysed in RIPA buffer with sonication or fixed in 4% PFA for immunohistochemistry. MN cultures were grown for 14dpf and treated overnight with DMSO as control or 25 µM Etoposide. Cells were then fixed in 4% PFA and immune stained. Mean intensity of MNs per cell was counted within an ellipsoid ROI covering most of the cell and were averaged per image and condition. n=3 technical replicates per condition, with minimum 3 field of view images per replica and 7-10 cells per image. For HEK cells, Mean intensity per cell was counted within an ellipsoid ROI covering most of the cells. n=3, with > 40 cells measured per condition across 5-7 images.

PLK1 inhibitor treatment: neurons were differentiated from iPCS-derived NPCs as described above and were treated with 1 µM BI2536 for 24 hours, 2 µM Etoposide for 1 hour, or with the equivalent volume of DMSO. Neurons were then fixed in 4% paraformaldehyde for 20 minutes at 37°C. Afterwards, neurons were permeabilized in 0.2% Triton X-100 for 10 minutes. Then, MNs were washed in tris-buffered saline containing 1% Tween 20 for 10 minutes for 3 times and subsequently blocked in Pierce Protein-free blocking buffer (ThermoFisher: 37572) for 1 hour at room temperature.

### Immunostainings

PLK1 staining: MNs were grown in multi well cell plates for 14dpm. Imunnostaining for PLK1 was done with polyclonal PLK1 (208G4) Rabbit mAb #4513 (Cell Signaling technology, 4513S). All cell nuclei were counted by ‘analayze particles’ plugin in FIJI with manual correction. PLK1 high signal cells were counted manually. High PLK1 signal was considered an intensity which had at least double of mean intensity value averaged across 25 random cells. Mean intensity per cell was counted within an ellipsoid ROI covering most of the cell. At DIV14: n=3 experiments. Total cell numbers and high PLK1 positive cells within 3-10 field of view images analyzed per condition, per experiment.

DNA damage foci: The primary antibodies α-MAP2 (Abcam: ab5392; 1:100,000) and α-γH2AX (Millipore: 05-636; 1:500) were diluted in Pierce Protein-free blocking buffer (ThermoFisher Scientific: 37572) and incubated over-night at 4 °C on a shaker. Samples were then rinsed in TBS for 10 minutes three times and the secondary antibodies Alexa Fluor-488 α-chicken (Invitrogen: A11039) and Alexa Fluor-568 anti-rabbit (Invitrogen: A11036) were diluted 1:5000 in TBS and incubated for 1 hour at room temperature, on a shaker. Samples were rinsed three times in TBS for 10 minutes and then mounted in DAPI Fluoromount-G mounting medium (Southern Biotechnology: 0100-20).

### Microscopy

Images were acquired on a Zeiss inverted AxioObserver.Z1 microscope with LSM 900 module and high-resolution Airyscan 2 module, using a 63x 1.4 NA plan apochromat objective. Data acquired from microscopy were evaluated using Fiji v1.53t

### Statistical analysis

All statistical analyses including Hierarchical clustering (with Euclidean distance measure and average linkage clustering), QC assessment, and cross-comparison studies were performed using R-studio software and *Partek™ Flow™* software, v11.0. Differential expression was analyzed by applying a adjusted *P* ≤ 0.05 to the DESeq2 results followed by false discovery rate (FDR) step-up for multiple comparisons. Genes with FDR/adjusted *p*-value ≤ 0.05 and log2FC ≥ 1 or log2FC ≤ −1 were considered as significant thresholds for the identification of DEGs. All of the gene expression samples presented in this study were designed for 3 biological replicates (mean ± standard error (SEM), n ≥ 3). For the functional enrichment analysis, significantly enriched GO terms, pathways and PPI networks were identified using a *p*-value ≤ 0.05 as the cut off value for statistical significance. GSEA was performed in *Partek™ Flow™* software, v11.0 using normalized gene count values. Validation data were plotted and statistically analyzed on GraphPad Prism v9.4.1. software. Bars represented the mean and standard error of the mean. Two-way ANOVA or Unpaired parametric Student’s *t*-test results were used to compare between groups. Differences were considered statistically significant for *p* values < 0.05. *p* values were depicted in graphs as *, **, ***, **** representing *p* ≤ 0.05, *p* ≤ 0.01, *p* ≤ 0.001, and *p* ≤ 0.0001, respectively.

## Author Contributions

Conceptualization, V.Z. B.D and A.H.; methodology, V.Z., B.D, D.G and A.H.; validation, V.Z., B.D.; formal analysis, V.Z., B.D and H.G.; investigation, V.Z. B.D., D.G., R.V., V.K., A.D., and H.G.; resources, A.H., S.R., E.Z., C.D., data curation, V.Z., B.D, and S.R.; writing—original draft preparation, V.Z., B.D, S.R., and A.H.; writing—review and editing, all authors; visualization, V.Z., B.D., and D.G., ; supervision, A.H.,., S.R., C.D., and E.Z.; project administration, A.H.; funding acquisition, A.H., S.R., C.D., and E.Z. All authors have read and agreed to the published version of the manuscript.

## Institutional Review Board Statement

The performed procedures were in accordance with the Declaration of Helsinki (WMA, 1964) and approved by the Ethical Committee of the Technische Universität Dresden, Germany (EK 393122012 and EK 45022009) Informed Consent Statement: Written informed consent was obtained from all participants, including for publication of any research results.

## Data and code availability

All sequencing data have been deposited to GEO under accession number GEO: GSE276214.

## Supporting information

Differential gene expression results from FUSP525L versus FUSWT iPSC-derived MNs on the axonal side

Differential gene expression results from FUSWT Axon versus FUSWT Soma iPSC-derived MNs

GSEA enrichment analysis (GO gene sets) results for normalized gene counts from FUSWT Axon versus FUSWT Soma iPSC-derived MNs

RNA-seq datasets used for cross comparison analysis

GSEA enrichment analysis (GO gene sets) results for normalized gene counts from FUSP525L versus FUSWT iPSC derived MNs enriched on the somatodendritic

PPI analysis results (Up DEGs) from FUSP525L versus FUSWT iPSC derived MNs on the axonal side (Parameter setting: interaction score 0.40

GO term analysis results of the Axon transcriptome (Up DEGs) from FUSP525L versus FUSWT iPSC-derived MNs

Cell lines used in this study

## Acknowledgments

We deeply thank the patients and partners who took part in this study. We acknowledge the great cell culture help of Jette Abel. This work was supported by next-generation sequencing facility (Dresden-concept Genome Center) for their excellent RNA-sequencing services and bioinformatics support, in particular, Mathias Lesche and Andreas Petzold. We would like to thank Wendy Bergmann and Michael Müller from the Core Facility for Cell Sorting and Cell Analysis at the Rostock University Medical Center for their kind assistance in the flow cytometric analyses.

