## Supplemental figures 1-12 for "Axonal transcriptome reveals upregulation of PLK1 as a protective mechanism in response to increased DNA damage in FUS^P525L^ spinal motor neurons"

Figure S1

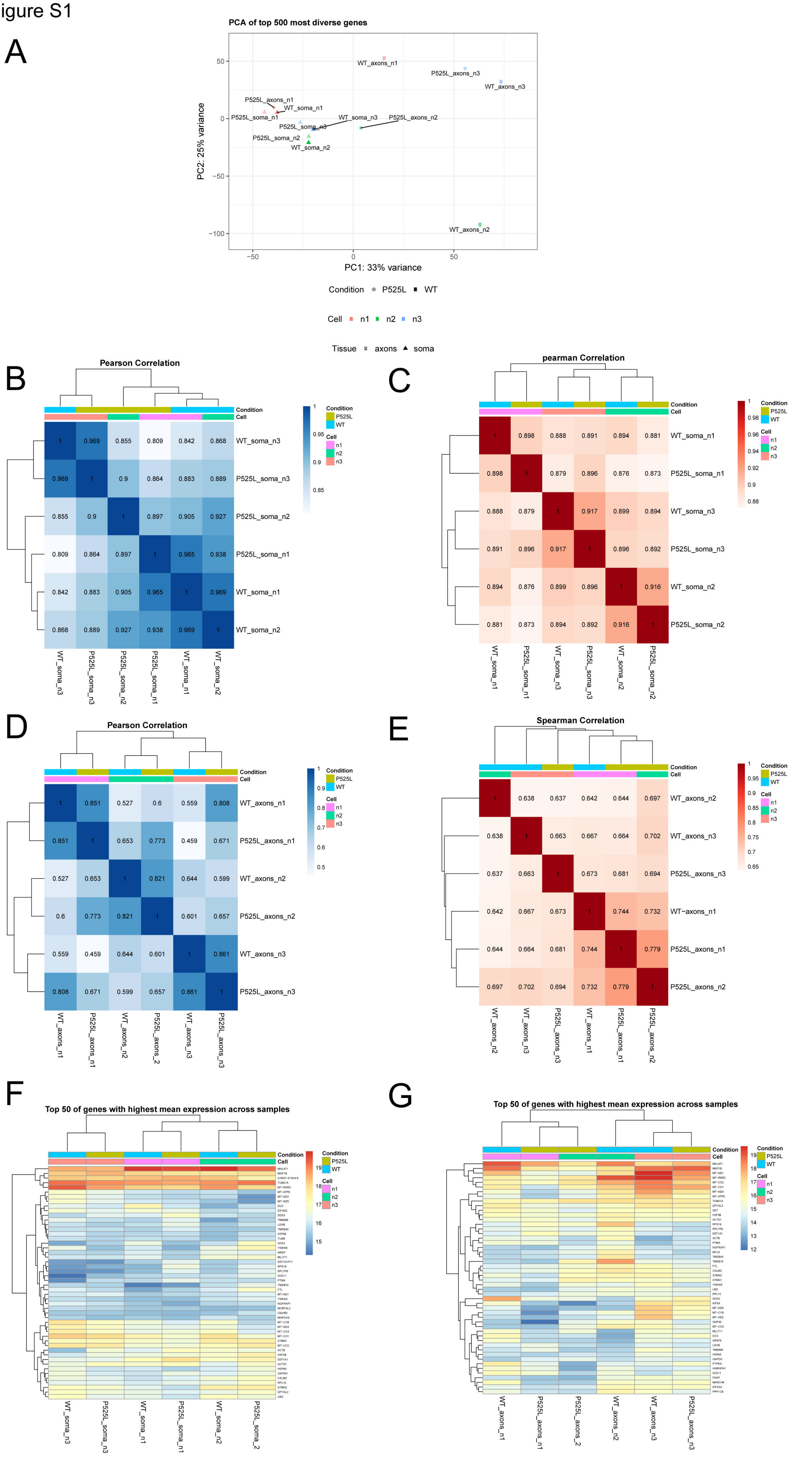
A
PCA of top 500 most diverse genes
PC1: 33% variance
PC2: 25% variance
WT_axons_n1
P525L_axons_n3
WT_axons_n3
P525L_axons_n1
WT_soma_n1
P525L_soma_n1
WT_soma_n3
P525L_axons_n2
P525L_soma_n3
P525L_soma_n2
WT_soma_n2
WT_axons_n2
Condition P525L WT
Cell n1 n2 n3
Tissue axons soma
B
Pearson Correlation
WT_soma_n3
P525L_soma_n3
P525L_soma_n2
P525L_soma_n1
WT_soma_n1
WT_soma_n2
C
pearman Correlation
WT_soma_n1
P525L_soma_n1
WT_soma_n3
P525L_soma_n3
WT_soma_n2
P525L_soma_n2
D
Pearson Correlation
WT_axons_n1
P525L_axons_n1
WT_axons_n2
P525L_axons_n2
WT_axons_n3
P525L_axons_n3
E
Spearman Correlation
WT_axons_n2
WT_axons_n3
P525L_axons_n3
WT-axons_n1
P525L_axons_n1
P525L_axons_n2
F
Top 50 of genes with highest mean expression across samples
WT_soma_n3
P525L_soma_n3
WT_soma_n1
P525L_soma_n1
WT_soma_n2
P525L_soma_2
G
Top 50 of genes with highest mean expression across samples
WT_axons_n1
P525L_axons_n1
P525L_axons_2
WT_axons_n2
WT_axons_n3
P525L_axons_n3
Condition
Cell

### Figure S2

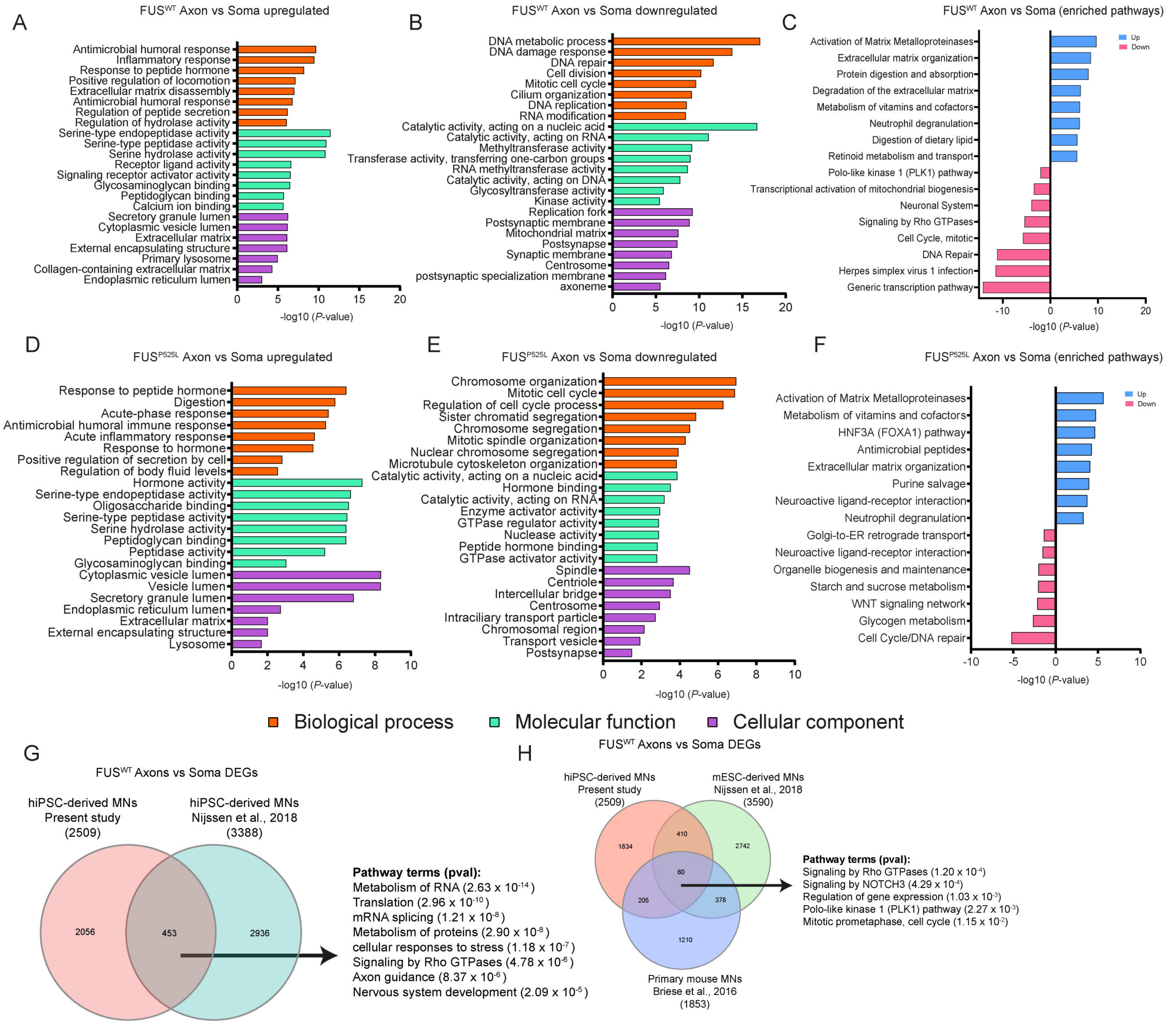
A
FUSWT Axon vs Soma upregulated
Antimicrobial humoral response
Inflammatory response
Response to peptide hormone
Positive regulation of locomotion
Extracellular matrix disassembly
Antimicrobial humoral response
Regulation of peptide secretion
Regulation of hydrolase activity
Serine-type endopeptidase activity
Serine-type peptidase activity
Serine hydrolase activity
Receptor ligand activity
Signaling receptor activator activity
Glycosaminoglycan binding
Peptidoglycan binding
Calcium ion binding
Secretory granule lumen
Cytoplasmic vesicle lumen
Extracellular matrix
External encapsulating structure
Primary lysosome
Collagen-containing extracellular matrix
Endoplasmic reticulum lumen
0 5 10 15 20
-log10 (P-value)
B
FUSWT Axon vs Soma downregulated
DNA metabolic process
DNA damage response
DNA repair
Cell division
Mitotic cell cycle
Cilium organization
DNA replication
RNA modification
Catalytic activity, acting on a nucleic acid
Catalytic activity, acting on RNA
Methyltransferase activity
Transferase activity, transferring one-carbon groups
RNA methyltransferase activity
Catalytic activity, acting on DNA
Glycosyltransferase activity
Kinase activity
Replication fork
Postsynaptic membrane
Mitochondrial matrix
Postsynapse
Synaptic membrane
Centrosome
postsynaptic specialization membrane
axoneme
0 5 10 15 20
-log10 (P-value)
C
FUSWT Axon vs Soma (enriched pathways)
Up
Down
Activation of Matrix Metalloproteinases
Extracellular matrix organization
Protein digestion and absorption
Degradation of the extracellular matrix
Metabolism of vitamins and cofactors
Neutrophil degranulation
Digestion of dietary lipid
Retinoid metabolism and transport
Polo-like kinase 1 (PLK1) pathway
Transcriptional activation of mitochondrial biogenesis
Neuronal System
Signaling by Rho GTPases
Cell Cycle, mitotic
DNA Repair
Herpes simplex virus 1 infection
Generic transcription pathway
-10 0 10 20
-log10 (P-value)
D
FUSP525L Axon vs Soma upregulated
Response to peptide hormone
Digestion
Acute-phase response
Antimicrobial humoral immune response
Acute inflammatory response
Response to hormone
Positive regulation of secretion by cell
Regulation of body fluid levels
Hormone activity
Serine-type endopeptidase activity
Oligosaccharide binding
Serine-type peptidase activity
Serine hydrolase activity
Peptidoglycan binding
Peptidase activity
Glycosaminoglycan binding
Cytoplasmic vesicle lumen
Vesicle lumen
Secretory granule lumen
Endoplasmic reticulum lumen
Extracellular matrix
External encapsulating structure
Lysosome
0 2 4 6 8 10
-log10 (P-value)
E
FUSP525L Axon vs Soma downregulated
Chromosome organization
Mitotic cell cycle
Regulation of cell cycle process
Sister chromatid segregation
Chromosome segregation
Mitotic spindle organization
Nuclear chromosome segregation
Microtubule cytoskeleton organization
Catalytic activity, acting on a nucleic acid
Hormone binding
Catalytic activity, acting on RNA
Enzyme activator activity
GTPase regulator activity
Nuclease activity
Peptide hormone binding
GTPase activator activity
Spindle
Centriole
Intercellular bridge
Centrosome
Intraciliary transport particle
Chromosomal region
Transport vesicle
Postsynapse
0 2 4 6 8 10
-log10 (P-value)
F
FUSP525L Axon vs Soma (enriched pathways)
Up
Down
Activation of Matrix Metalloproteinases
Metabolism of vitamins and cofactors
HNF3A (FOXA1) pathway
Antimicrobial peptides
Extracellular matrix organization
Purine salvage
Neuroactive ligand-receptor interaction
Neutrophil degranulation
Golgi-to-ER retrograde transport
Neuroactive ligand-receptor interaction
Organelle biogenesis and maintenance
Starch and sucrose metabolism
WNT signaling network
Glycogen metabolism
Cell Cycle/DNA repair
-10 -5 0 5 10
-log10 (P-value)
Biological process
Molecular function
Cellular component
G
FUSWT Axons vs Soma DEGs
hiPSC-derived MNs
Present study
(2509)
hiPSC-derived MNs
Nijssen et al., 2018
(3388)
2056
453
2936
Pathway terms (pval):
Metabolism of RNA (2.63 x 10-14)
Translation (2.96 x 10-10)
mRNA splicing (1.21 x 10-8)
Metabolism of proteins (2.90 x 10-8)
cellular responses to stress (1.18 x 10-7)
Signaling by Rho GTPases (4.78 x 10-6)
Axon guidance (8.37 x 10-6)
Nervous system development (2.09 x 10-5)
H
FUSWT Axons vs Soma DEGs
hiPSC-derived MNs
Present study
(2509)
mESC-derived MNs
Nijssen et al., 2018
(3590)
1834
410
2742
60
205
378
1210
Primary mouse MNs
Briese et al., 2016
(1853)
Pathway terms (pval):
Signaling by Rho GTPases (1.20 x 10-4)
Signaling by NOTCH3 (4.29 x 10-4)
Regulation of gene expression (1.03 x 10-3)
Polo-like kinase 1 (PLK1) pathway (2.27 x 10-3)
Mitotic prometaphase, cell cycle (1.15 x 10-2)

Figure S3

### GSEA enrichment in $FUS^{WT}$ Axon versus Soma

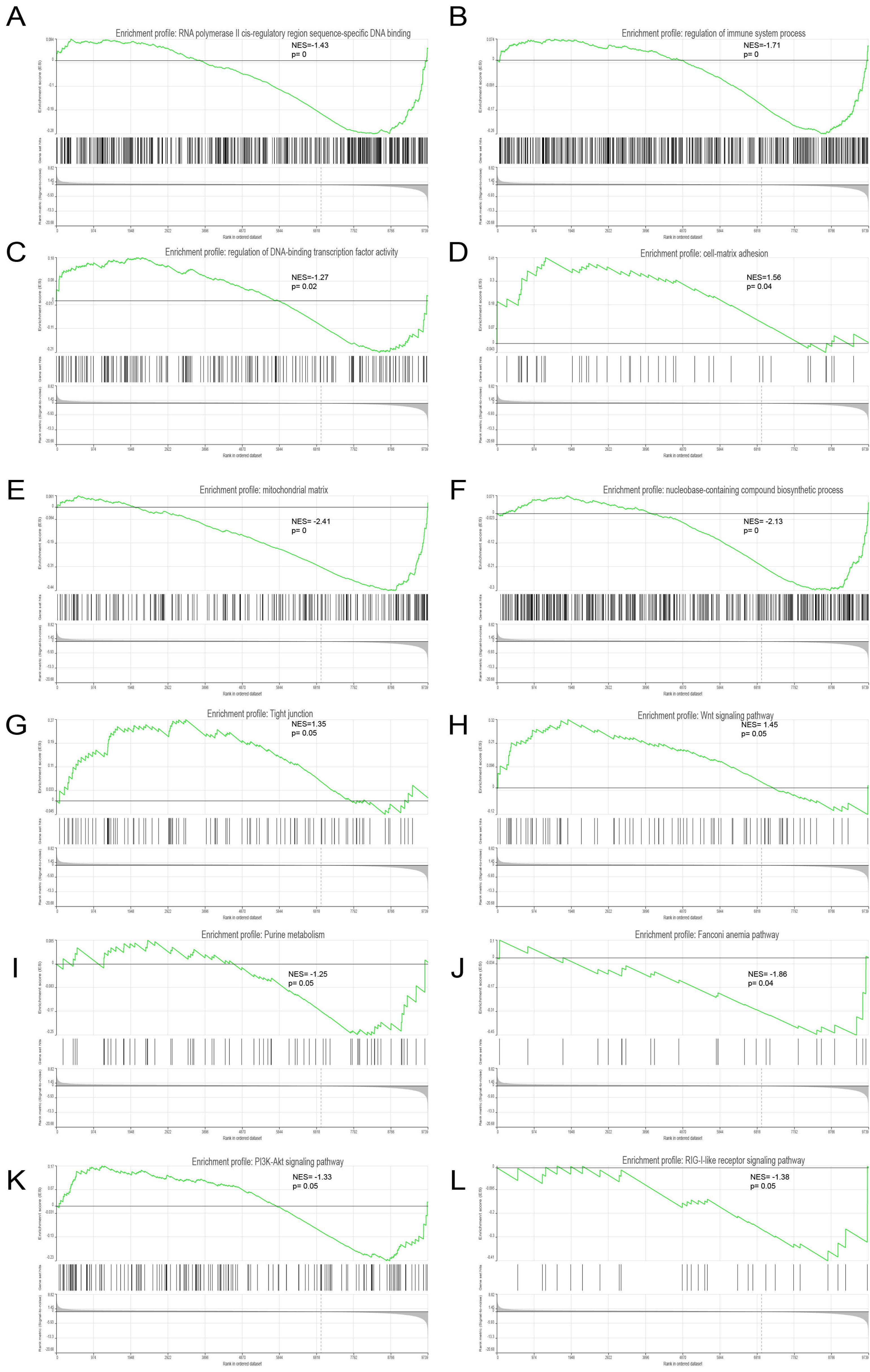

Figure S4

### GSEA enrichment in FUP$^{525L}$ Axon versus Soma

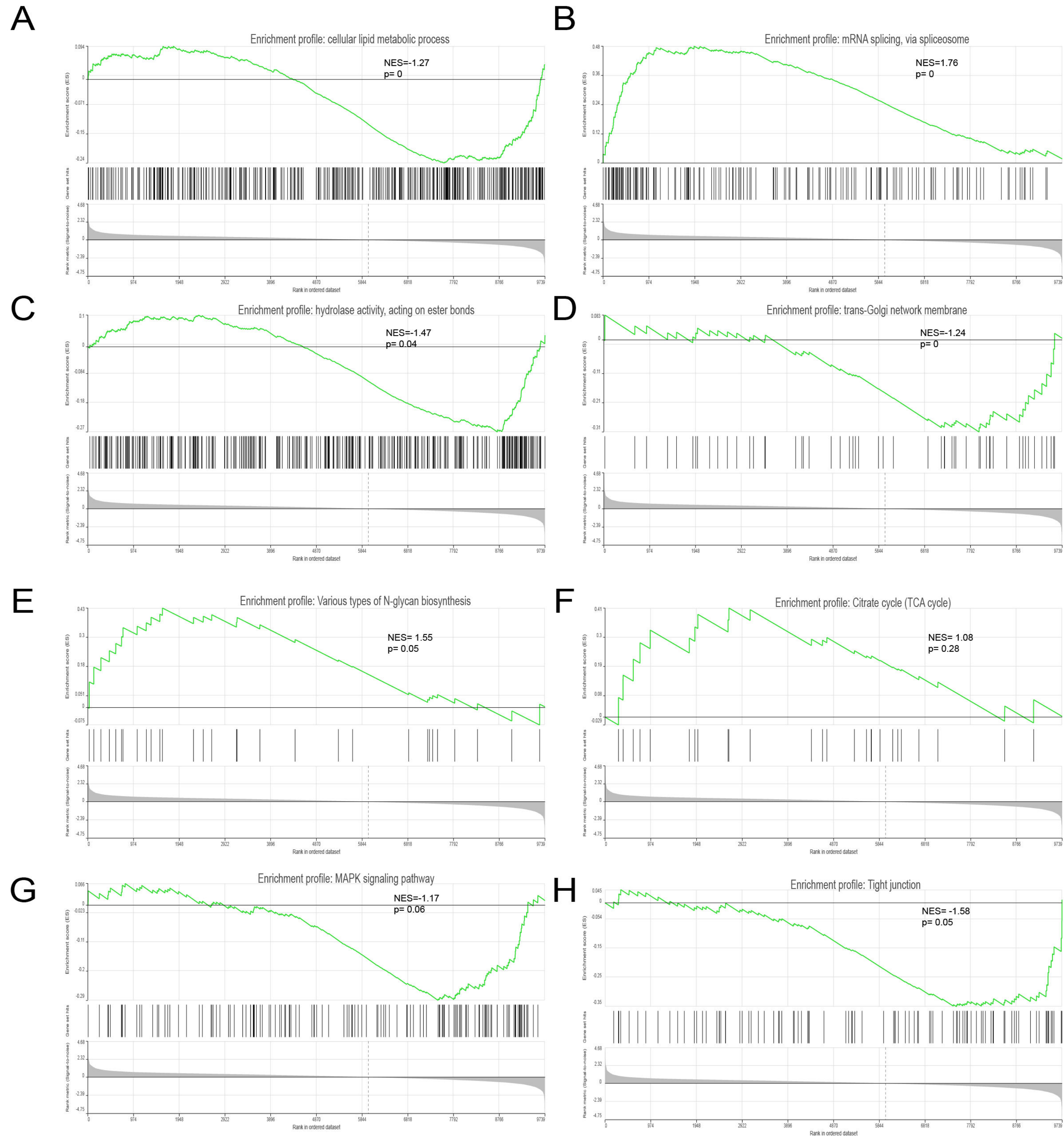

Figure S5

### GSEA enrichment in FUS$^{P525L}$ versus FUS$^{WT}$ Soma

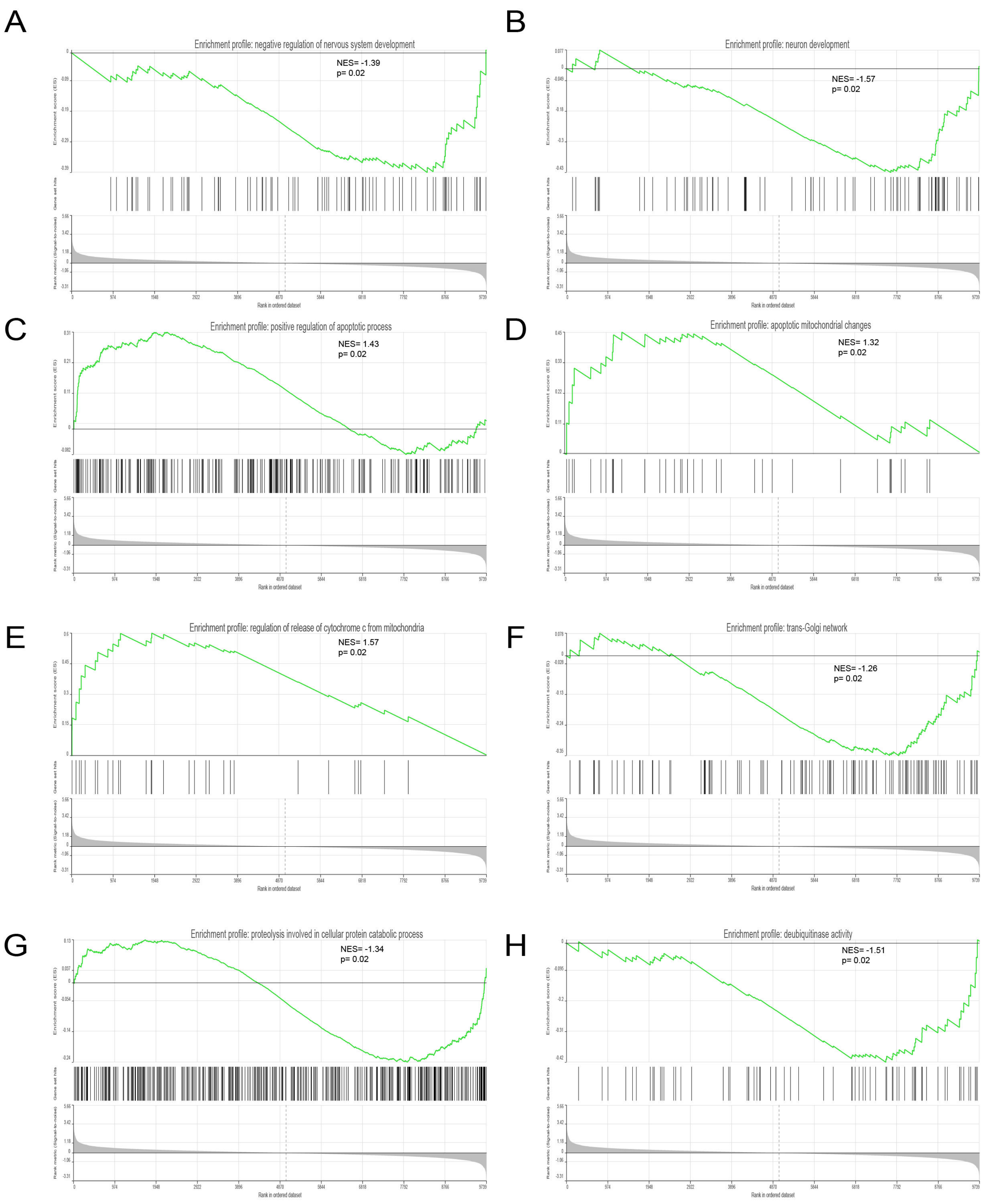

Figure S6

### GSEA enrichment in $FUS^{P525L}$ versus $FUS^{WT}$ Axon

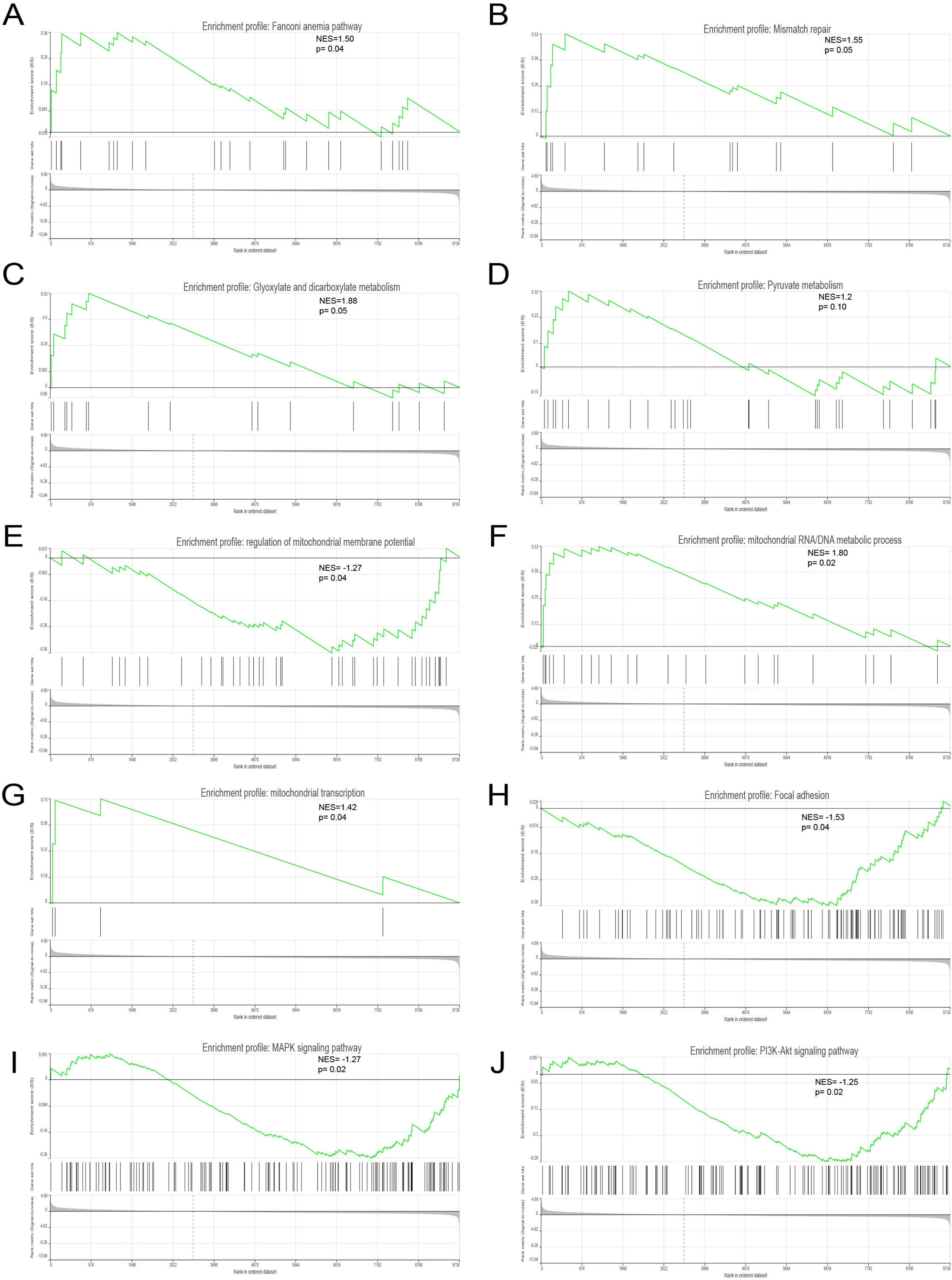

### Figure S7

#### A Cell cycle profiles for NPCs treated for 24 hours at different concentrations of BI2536

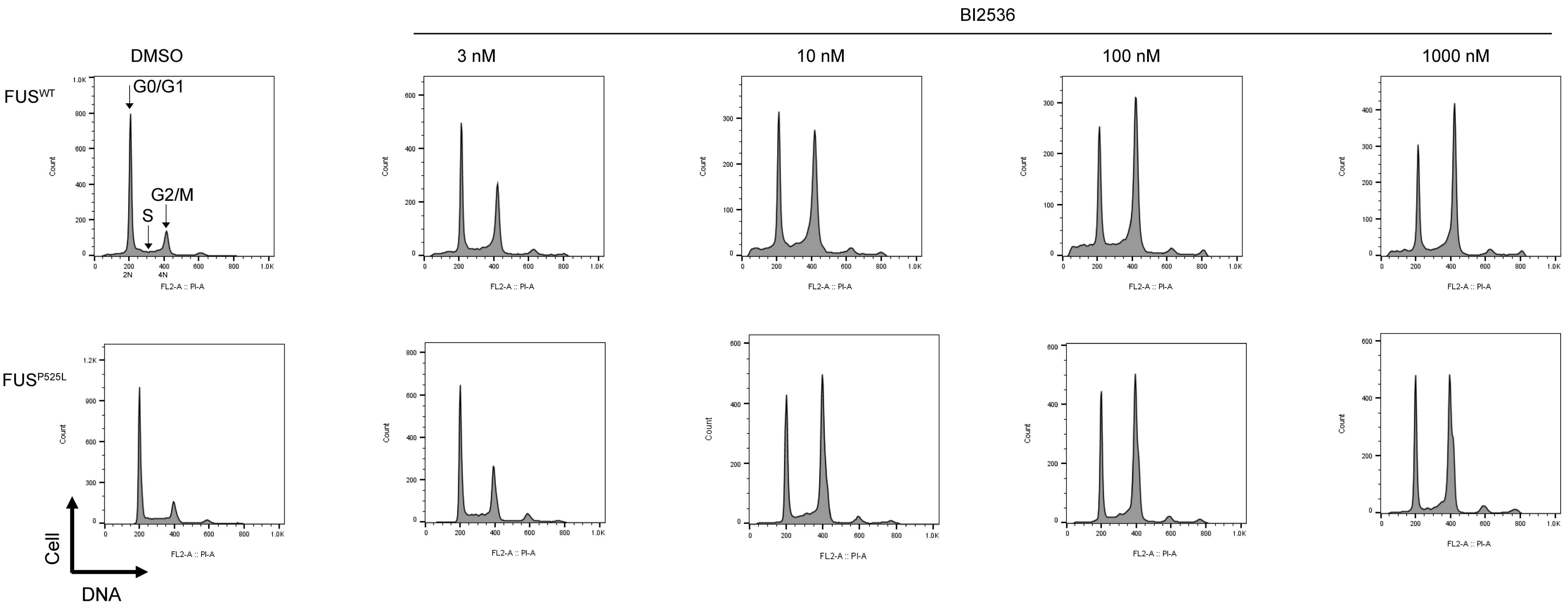


#### B Flow cytometric gating analysis of DNA cell cycle after PI staining in FUSWT NPCs (untreated)

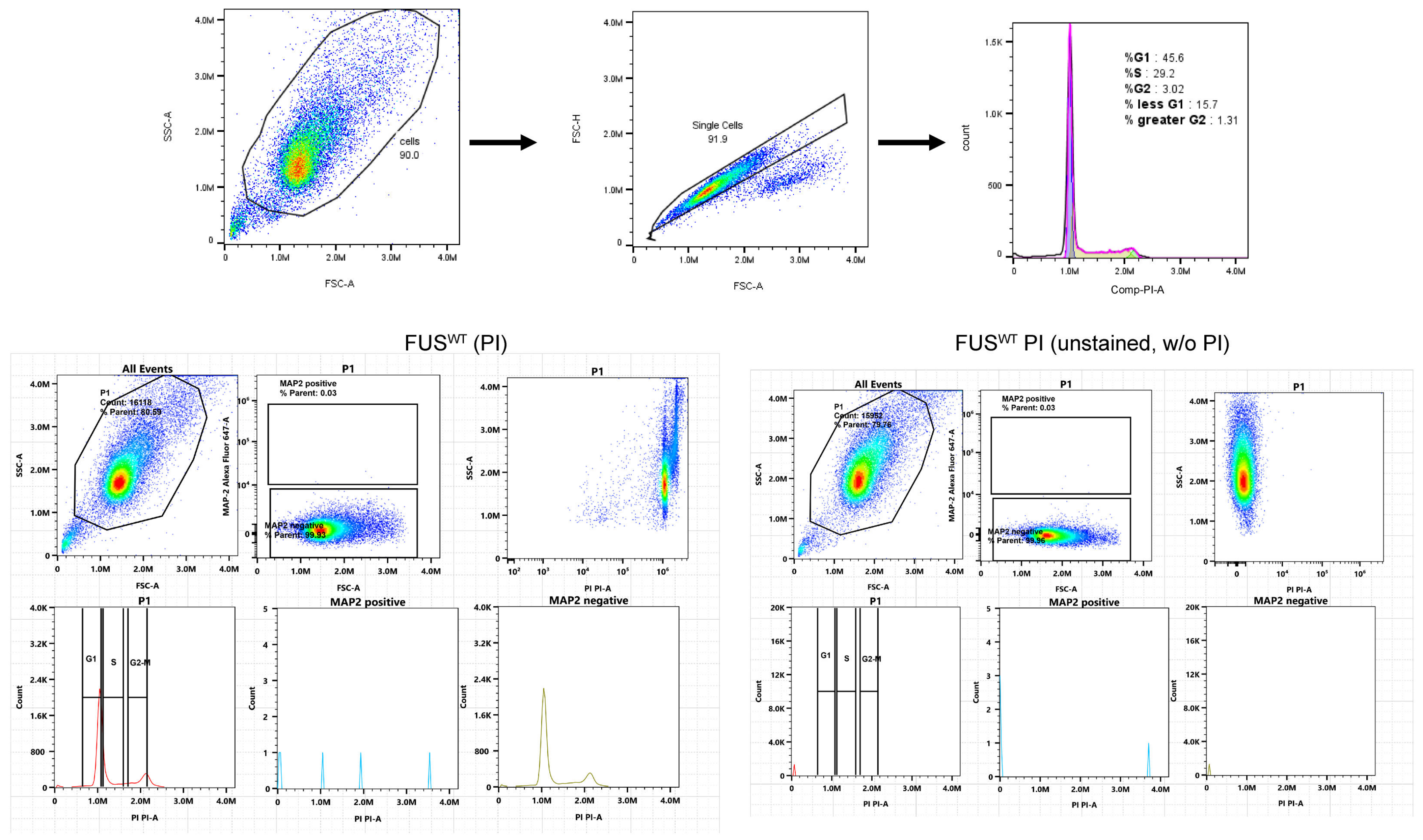

Figure S8

Flow cytometric gating analysis of DNA cell cycle after PI staining in $FUS^{WT}$ MAP2+ MNs (untreated)

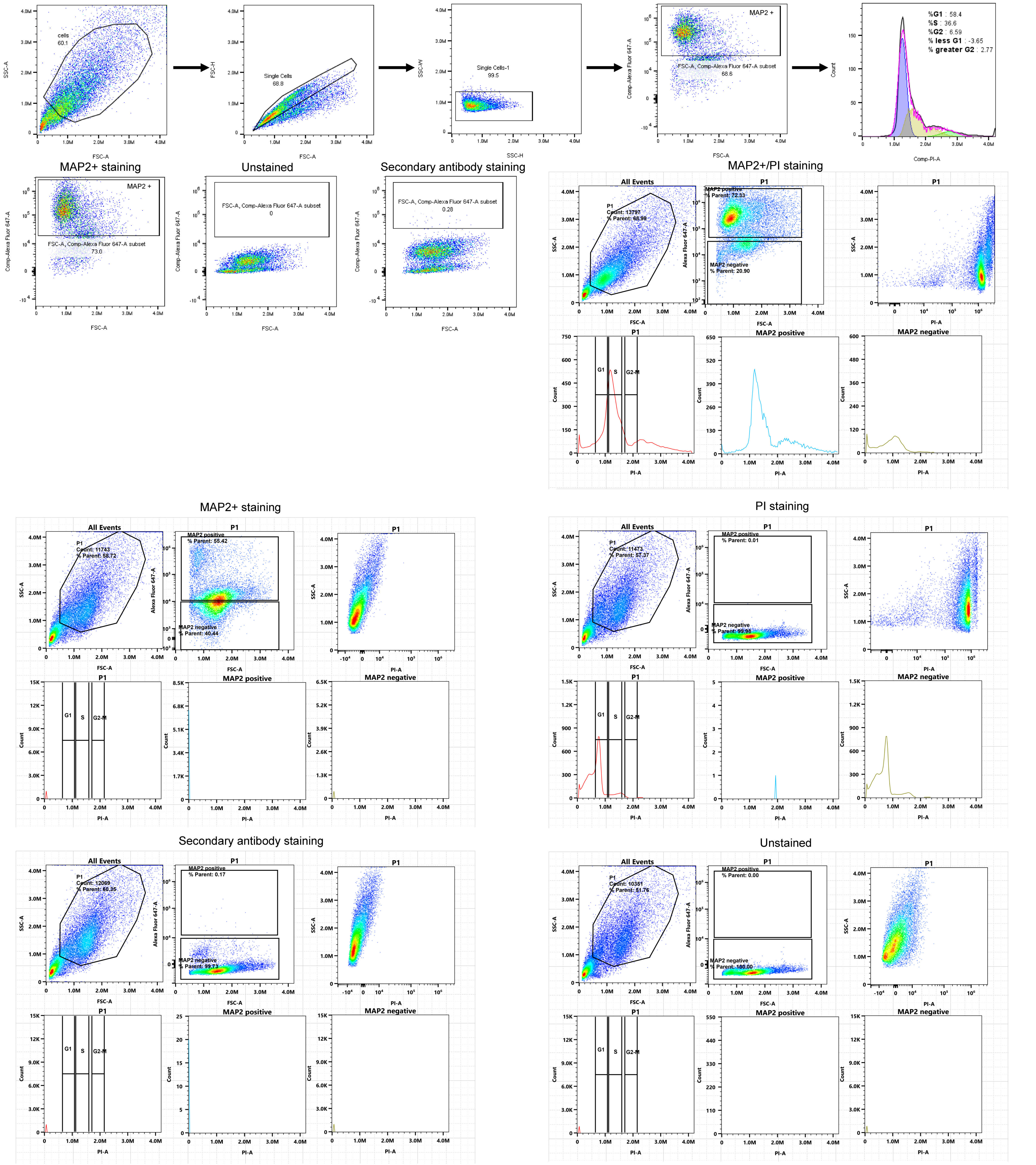

Figure S9

#### Flow cytometric gating analysis of DNA cell cycle after PI staining in FUS$^{P525L}$ MAP2+ MNs (untreated)

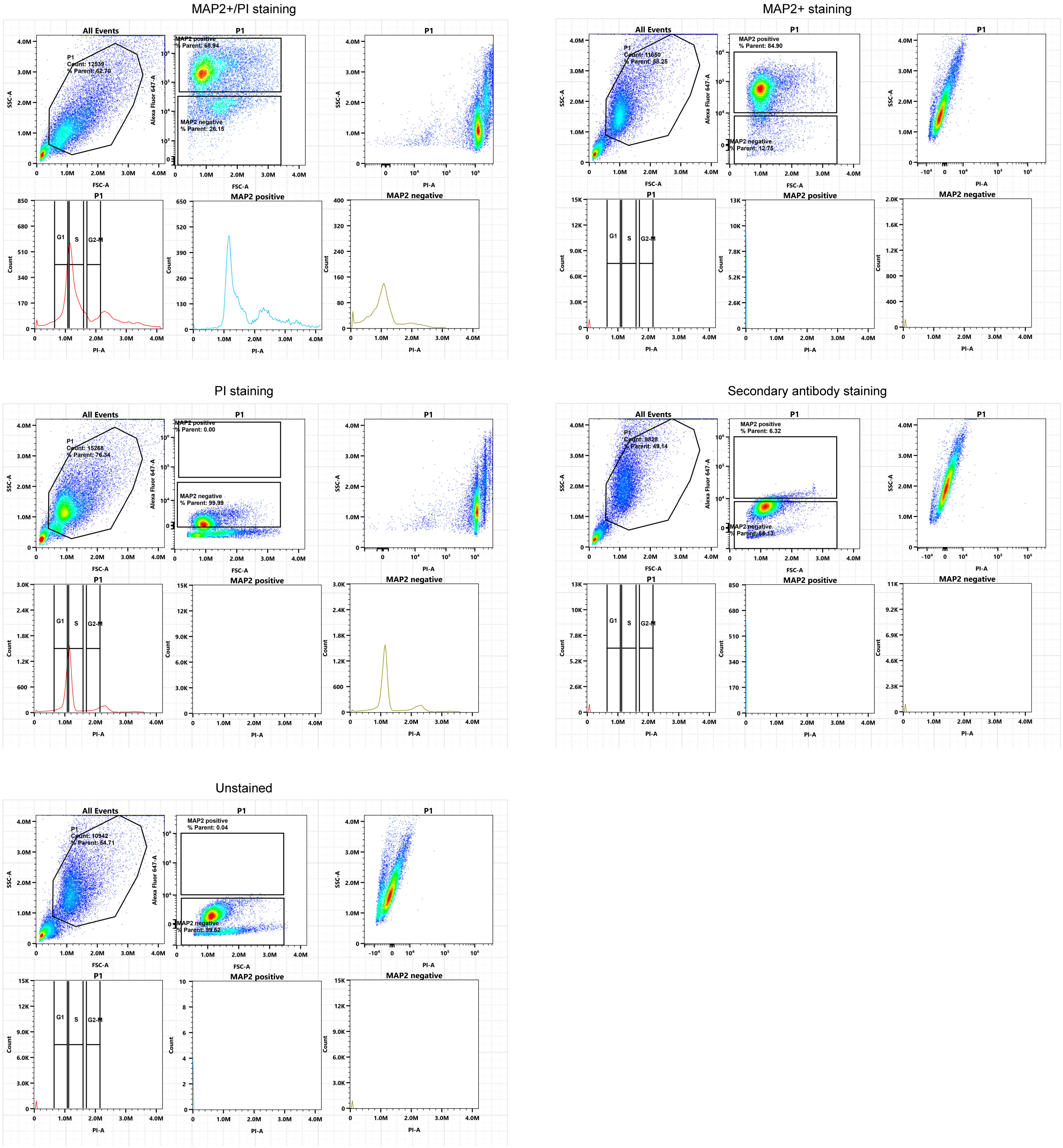

Figure S10

Annexin V-APC/PI multicolor staining assay to evaluate cell necrosis and apoptosis in $FUS^{WT}$ NeuO+MNs

A

DMSO

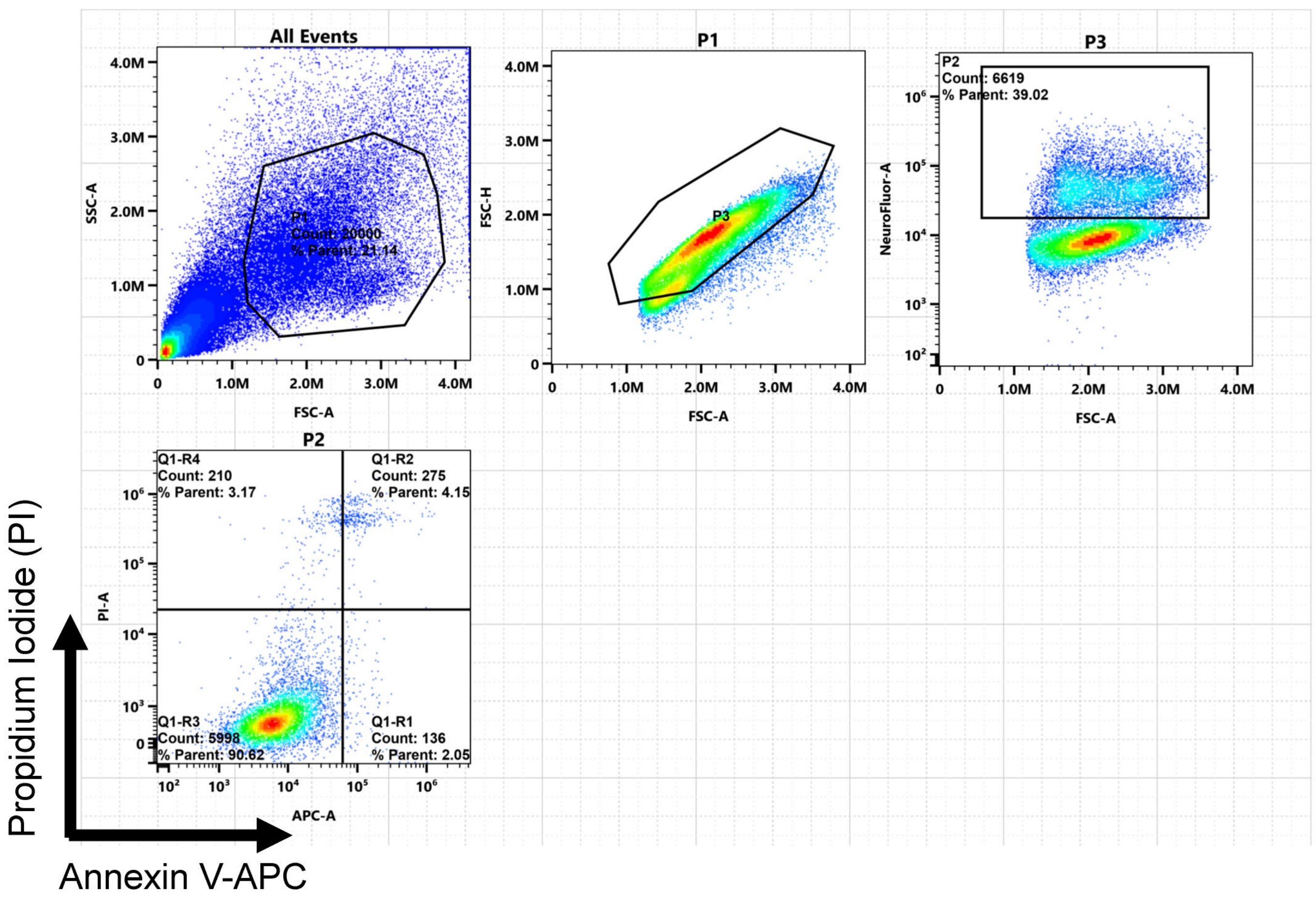


B

BI2536 100 nM

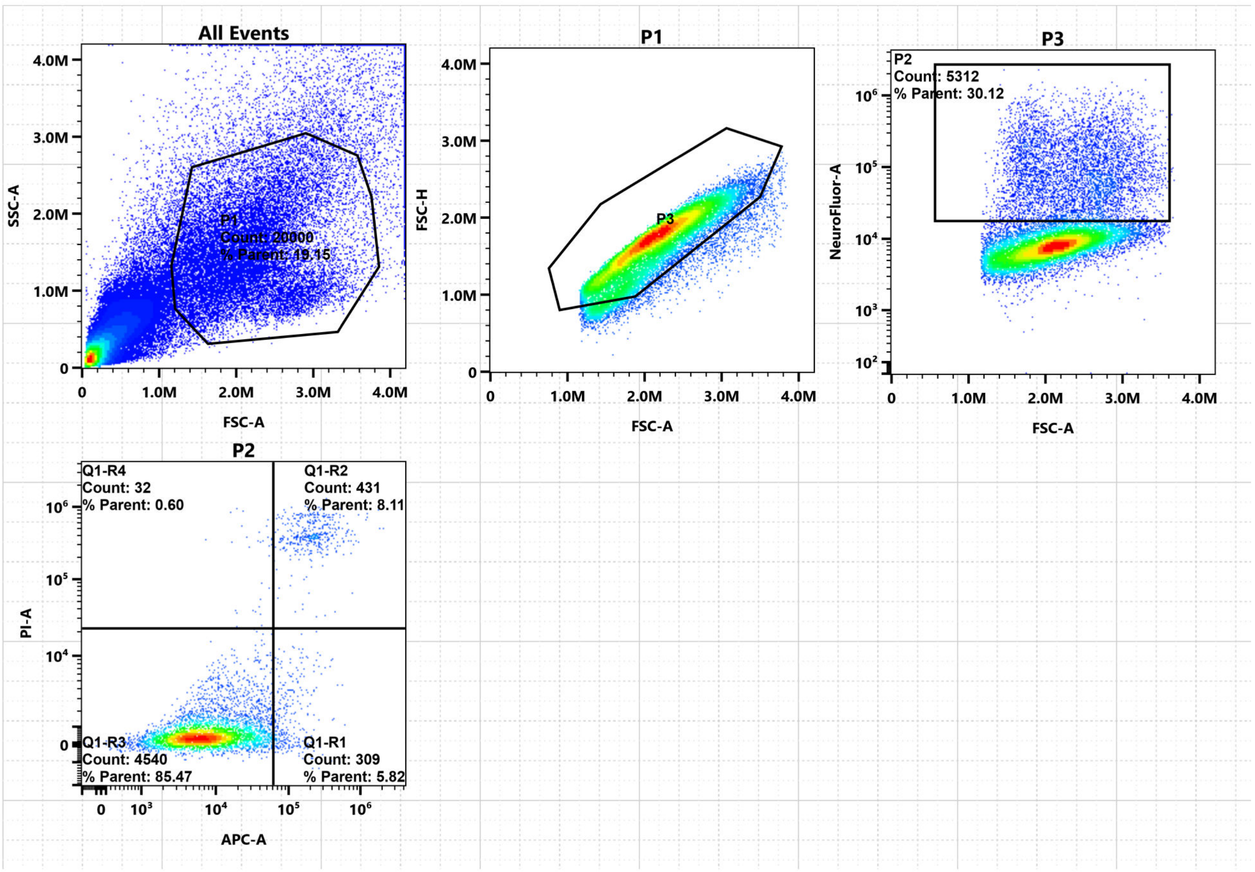


C

Unstained

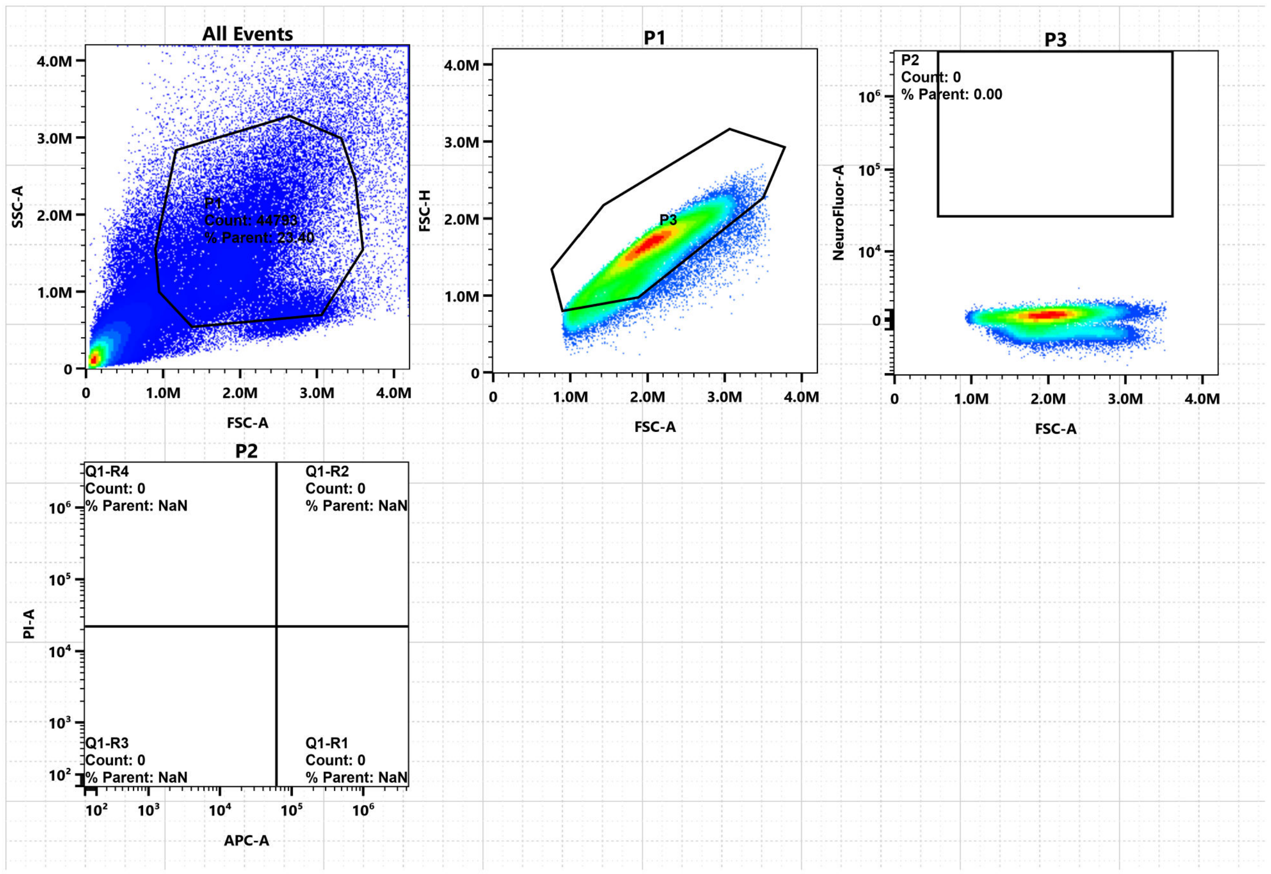


D

NeuO+ staining

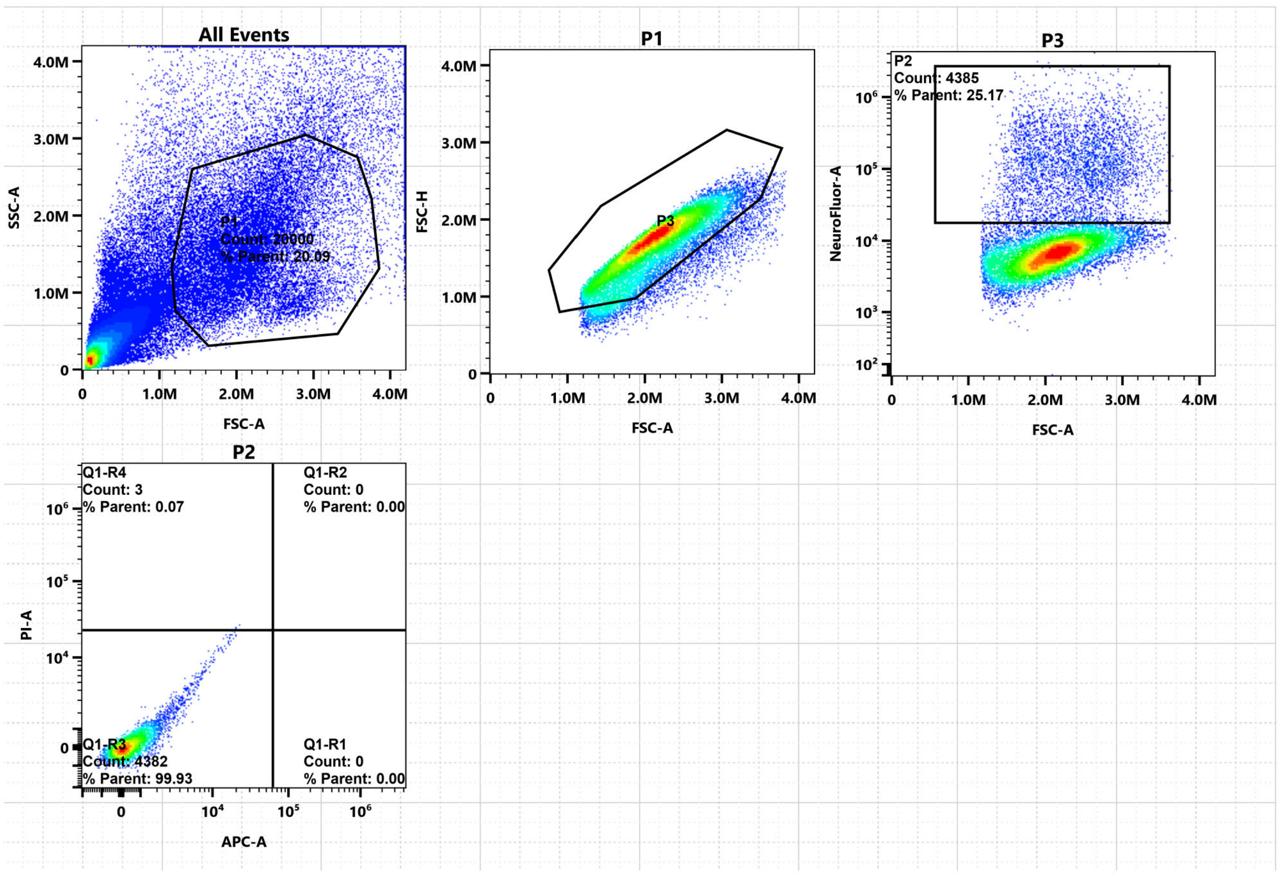

### Figure S11

Annexin V-APC/PI multicolor staining assay to evaluate cell necrosis and apoptosis in $FUS^{P525L}$ NeuO+ MNs

A

DMSO

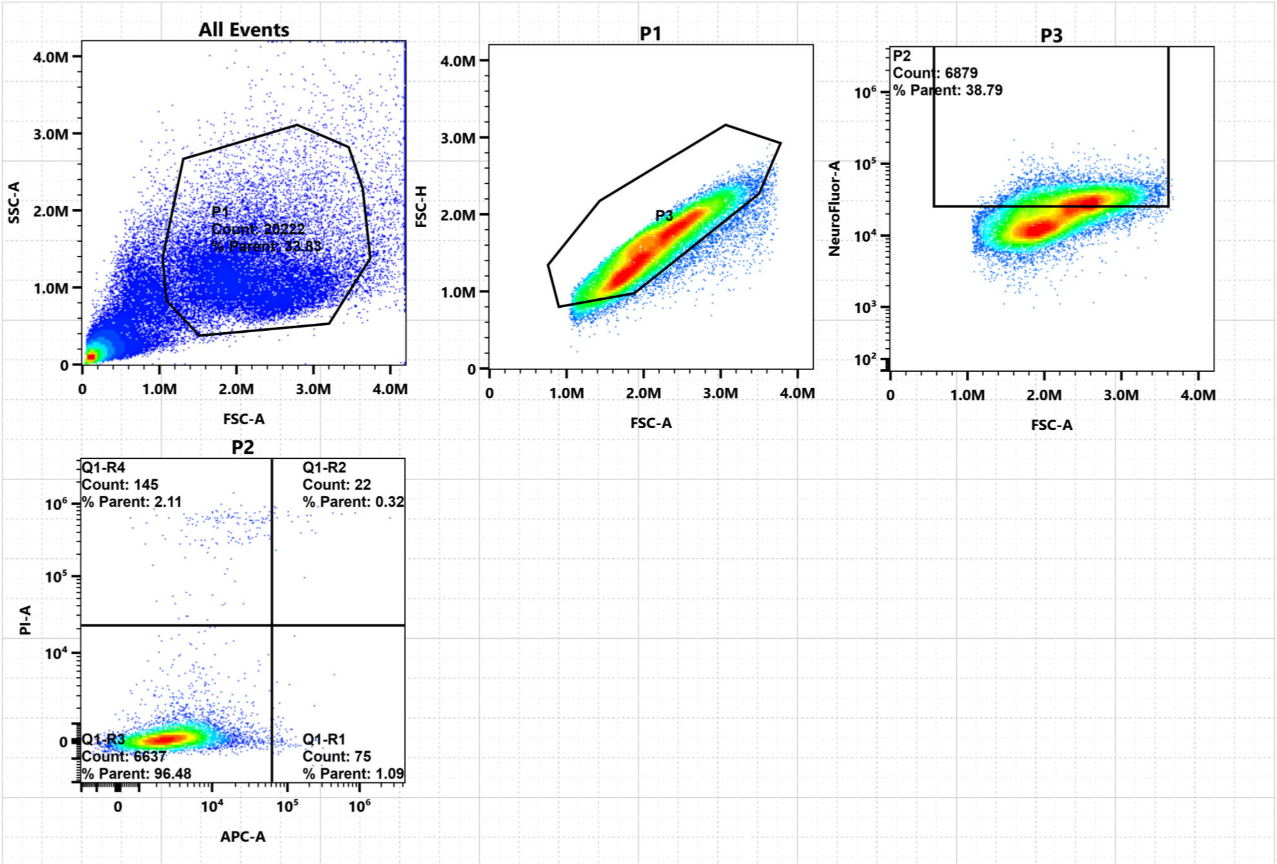


B

BI2536 100 nM

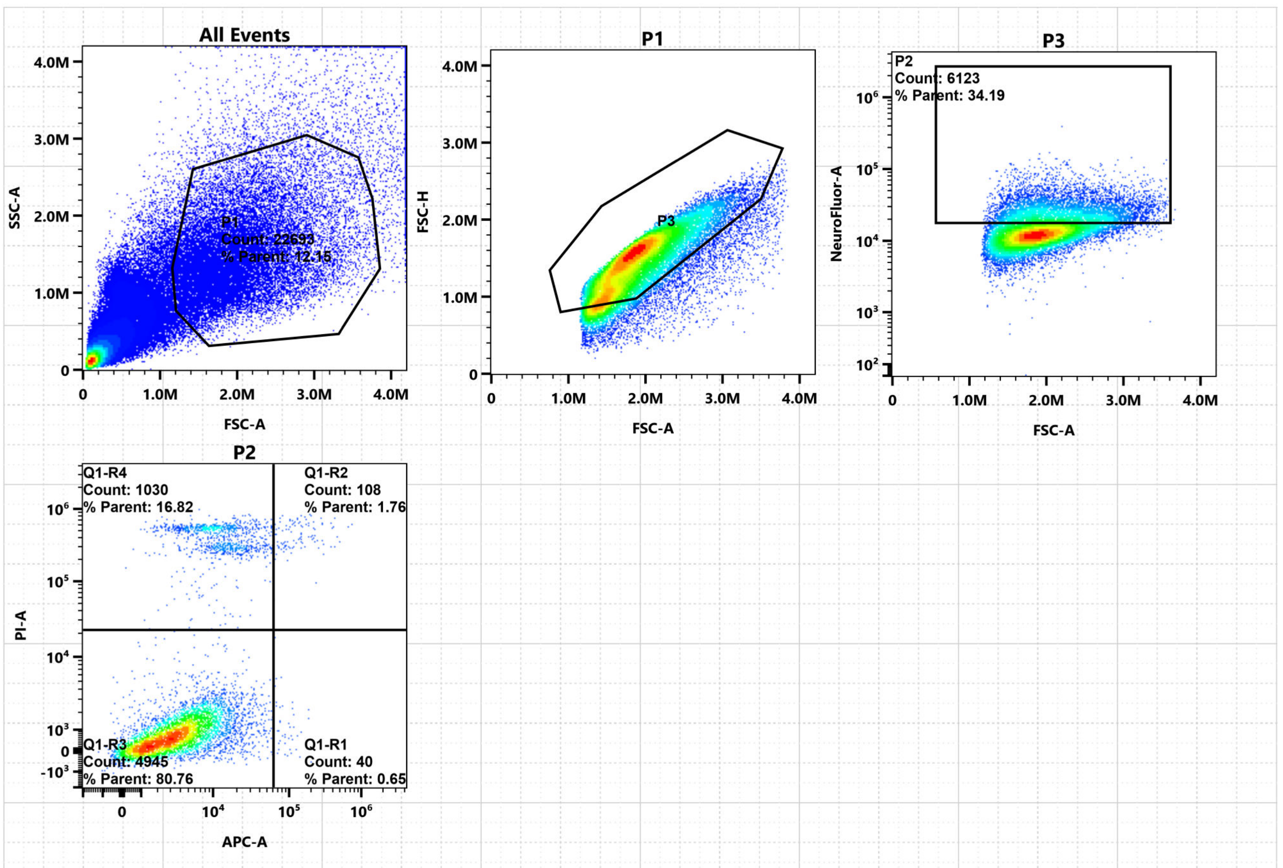


C

Unstained

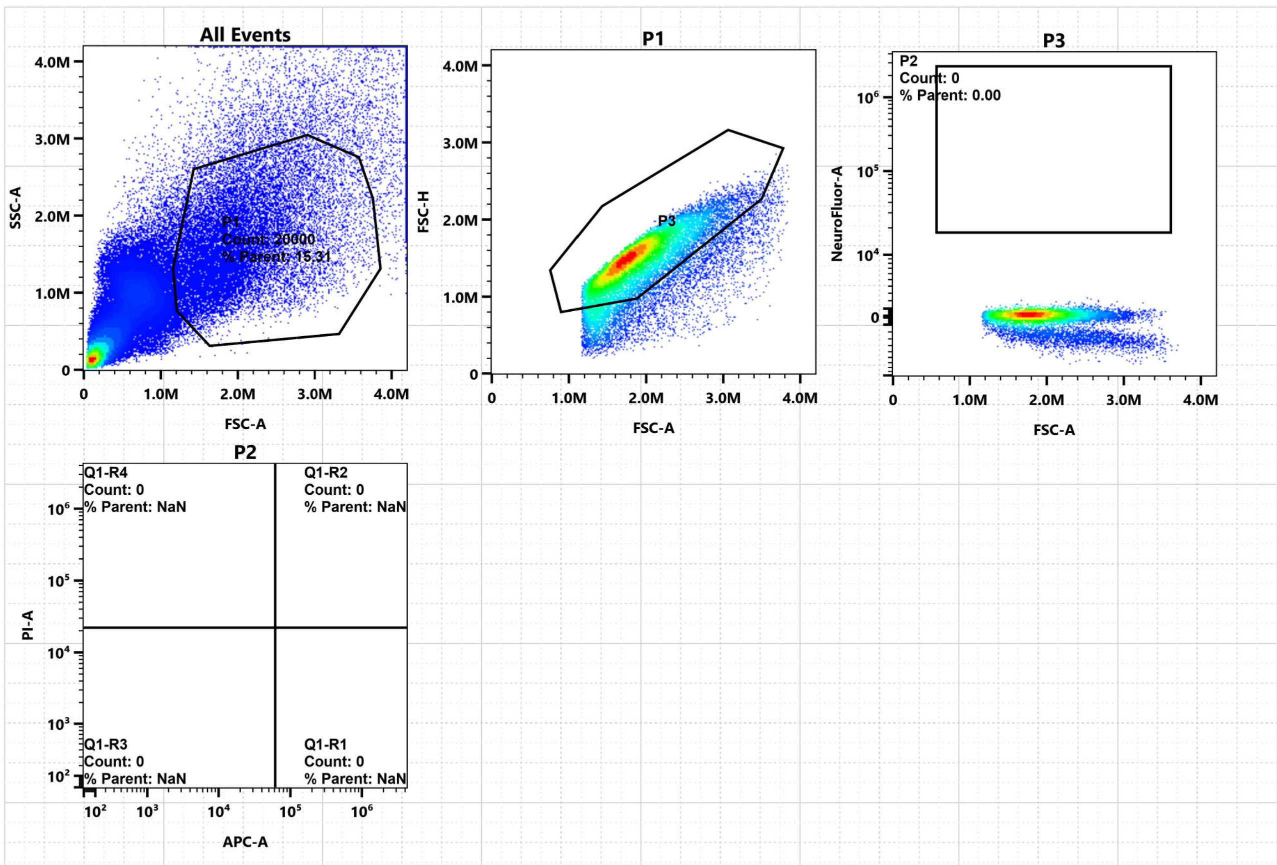


D

NeuO+ staining

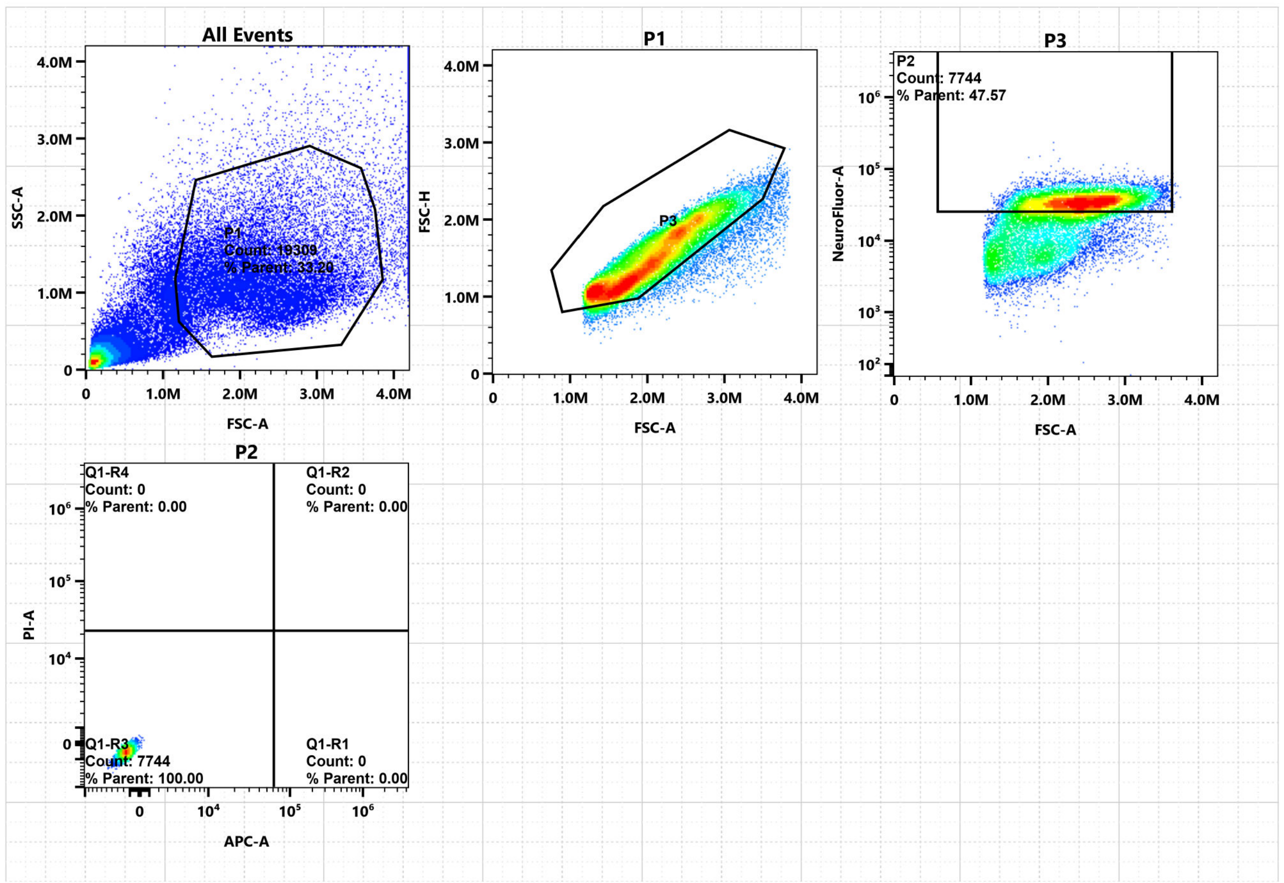

Figure S12

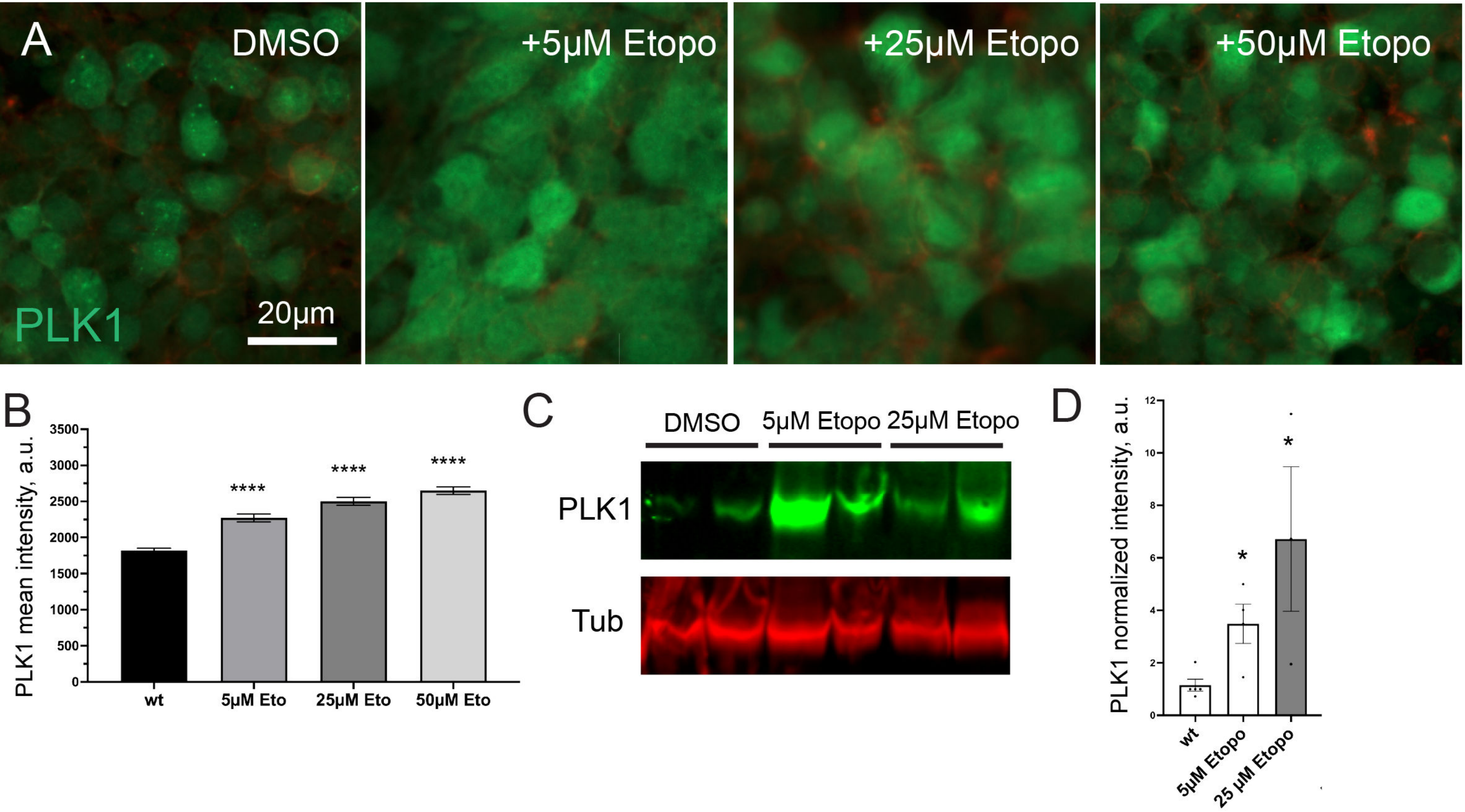

A
DMSO
+5µM Etopo
+25µM Etopo
+50µM Etopo
PLK1
20µm
B
PLK1 mean intensity, a.u.
3500
3000
2500
2000
1500
1000
500
0
****
****
****
wt
5µM Eto
25µM Eto
50µM Eto
C
DMSO
5µM Etopo
25µM Etopo
PLK1
Tub
D
PLK1 normalized intensity, a.u.
12
10
8
6
4
2
0
*
*
wt
5µM Etopo
25 µM Etopo
