## Supplementary material for "Axonal transcriptome reveals upregulation of PLK1 as a protective mechanism in response to increased DNA damage in FUS^P525L^ spinal motor neurons": Cell lines used in this study

**Supplementary Table S10:** Cell lines used in this study.

| **Genotype** | **Cell line** | **Sex** | **Age at biopsy** | **Mutation** | **Primarily characterized in** |
| --- | --- | --- | --- | --- | --- |
| Isogenic pair control1 | FUS^WT^-eGFP^het^ | isogenic to FUS ^R521C^ and FUS^P525L^ eGFP | N/A | - | (Naumann, Pal et al. 2018) |
| Isogenic pair Mut1 | FUS^P525L^-eGFP^het^ FUS ^R521C het^ | isogenic to FUS ^R521C^ and FUS^WT^ eGFP female | N/A | P525L | (Japtok, Lojewski et al. 2015, Naumann, Pal et al. 2018) |
| Control 2 | 2062-2 | male | 43 |  | (Peter, Trilck et al. 2017) |
| Control 3 | GM23251-4 | female | 41 |  | (Petters, Völkner et al. 2020) |
| Control 4 | AKC26 | female | 48 |  | (Grossmann, Malburg et al. 2023) |
| Mut2 | FUS^R521C^ | female | 58 | R521C | (Japtok, Lojewski et al. 2015, Naumann, Pal et al. 2018) |
| Mut3 | R495QfsX527 | male | 46 |  | (Japtok, Lojewski et al. 2015, Naumann, Pal et al. 2018) |
